# Chm7 and Heh1 form a nuclear envelope subdomain for nuclear pore complex quality control

**DOI:** 10.1101/049148

**Authors:** Brant M. Webster, David J. Thaller, Jens Jäeger, Sarah E. Ochmann, C. Patrick Lusk

## Abstract

Mechanisms that ensure the integrity of the nuclear envelope rely on membrane remodeling proteins like the ESCRTs and the AAA ATPase Vps4, which help seal the nuclear envelope at the end of mitosis and prevent the formation of defective nuclear pore complexes (NPCs). Here, we show that the integral inner nuclear membrane proteins Heh1 and Heh2 directly bind the ESCRT-III, Snf7, and the ESCRT-II/III chimera, Chm7, in their ‘open’ forms. Moreover, Heh1 is required for Chm7-recruitment to the nuclear envelope. As Chm7 accumulates on the nuclear envelope upon blocks to NPC assembly, but not to nuclear transport, interactions between ESCRTs and the Heh proteins might form a biochemically distinct nuclear envelope subdomain that delimits regions of assembling NPCs. Interestingly, deletion of *CHM7* suppresses the formation of the storage of improperly assembled NPC compartment prevalent in *vps4Δ* strains. Thus, our data support that the Heh1-dependent recruitment of Chm7 is a key component of a quality control pathway whose local regulation by Vps4 and the transmembrane nup, Pom152, prevents loss of nuclear compartmentalization by defective NPCs.

## Introduction

It is well established that the nuclear envelope (NE) in multicellular eukaryotes undergoes a dramatic breakdown and reformation during cell division (Wandke and Kutay, 2013), and it is emerging that the two membranes of the NE undergo extensive remodeling during interphase as well (Hatch and Hetzer, 2014; King and Lusk, 2016). Classic examples are the insertion of massive protein assemblies like nuclear pore complexes (NPCs)(Rothballer and Kutay, 2013) and the yeast centrosome (spindle pole body; SPB), but now extend to the nuclear egress of ‘mega’ ribonucleoprotein particles through a vesicular intermediate in the NE lumen/perinuclear space (Speese *et al.*, 2012; Jokhi *et al.*, 2013) and the degradative clearance of nuclear/NE contents by autophagy pathways that act specifically at the NE (Roberts *et al.*, 2003; Dou *et al.*, 2015; Mochida *et al.*, 2015). Understanding the molecular mechanisms that drive the membrane remodeling necessary for NE homeostasis is a critical goal for the field, particularly with the ever-growing links between disruptions in nuclear compartmentalization and human disease (Hatch and Hetzer, 2014; Burke and Stewart, 2014).

To exemplify the necessity of defining the molecular machineries capable of remodeling the NE, there is a lack of clarity regarding the fundamental mechanism of *de novo* NPC assembly. To build the massive ~50 MD yeast or ~100 MD human NPC requires the assembly of ~30 proteins (nucleoporins/nups) in multiple copies such that an individual NPC requires upwards of 500 proteins in yeast (Alber *et al.*, 2007) and perhaps twice as many in human cells (Bui *et al.*, 2013; von Appen *et al.*, 2015). This requires remarkable spatiotemporal control over hundreds of proteins that converge at a NE domain competent for NPC assembly; what defines a biogenesis site remains unclear but it might require local changes in Ran-GTP levels (Ryan and Wente, 2003; Walther *et al.*, 2003; D’Angelo *et al.*, 2006), nup binding to integral inner nuclear membrane (INM) proteins (Talamas and Hetzer, 2011; Yewdell *et al.*, 2011), or to chromatin (Franz *et al.*, 2007; Rasala *et al.*, 2008; Rotem *et al.*, 2009; Doucet *et al*., 2010), or changes in the properties of the membrane itself, perhaps by altering lipid composition (Schneiter *et al.*,1996; Scarcelli *et al*., 2007; Hodge *et al.*, 2010; Lone *et al.*, 2015). Local changes in lipid composition might also facilitate INM and outer nuclear membrane (ONM) fusion, but it is generally thought that protein-mediated membrane remodeling would ultimately be required to form a pore.

As yet, no single dedicated membrane bending or fusion machinery has been identified that drives nuclear pore formation but an emerging theme is the cooperative action of nups containing amphipathic helices or reticulon domains capable of recognizing or generating membrane curvature (Marelli *et al.*, 2001; Drin *et al.*, 2007; Dawson *et al*., 2009; Doucet *et al.*, 2010; Chadrin *et al*., 2010; Vollmer *et al.*, 2012; 2015; von Appen *et al.*, 2015; Mészáros *et al.*, 2015; Floch *et al.*, 2015; Casey *et al.*, 2015). Interestingly, budding yeast require the membrane bending and scission Endosomal Sorting Complexes Required for Transport (ESCRT)-III proteins to ensure formation of functional NPCs, raising the possibility that an established multifunctional membrane remodeler might yet be required during pore biogenesis (Webster *et al.*, 2014; Webster and Lusk, 2015). Similarly, recent work from *C. elegans* suggests that the ER lumenal AAA+ torsin orthologue OOC-5 might contribute to NPC assembly (VanGompel *et al.*, 2015). The concept that torsin, or one of its substrates, might be able to remodel membranes is supported by its role in the fission of INM evaginations during intralumenal bud formation necessary for mega-RNP egress (Jokhi *et al.*, 2013), and likely other yet to be defined processes (Goodchild *et al.*, 2005; Rose and Schlieker, 2012).

The membrane scission mechanism requiring torsins is topologically similar to that carried out by the ESCRTs; ESCRTs have been implicated in an ever-growing list of cellular processes that require a membrane scission step (Hurley, 2015). While the precise mechanism of membrane scission remains to be fully understood, in all cases it is thought that it will require the remarkable capacity of ESCRT-III’s to form polymers that generate and stabilize negative membrane curvature as membrane scaffolds (Ghazi-Tabatabai *et al.*, 2008; Lata, *et al.*, 2008; Hanson *et al.*, 2008; Saksena *et al.*, 2009; Henne *et al.*, 2012; Shen *et al.*, 2014; Cashikar *et al.*, 2014; Chiaruttini *et al.*, 2015).

There are multiple ESCRT-III proteins in budding yeast and multicellular eukaryotes with yeast Snf7 and its mammalian orthologue CHMP4B being the most abundant (Teis *et al.*, 2008); ESCRT-III’s are thought to share a similar structure with a basic core domain of four alpha helices whose assembly into a homo or hetero-polymer is autoinhibited by acidic helices that fold back onto the core (Muzioł *et al.*, 2006; Zamborlini *et al.*, 2006; Shim *et al.*, 2007; Kieffer *et al.*, 2008; Lata *et al.*, 2008; Xiao *et al.*, 2009; Bajorek *et al.*, 2009; Tang *et al.*, 2015). A key question therefore is how ESCRT-III’s come together to form potentially unique filaments with context-dependent biophysical properties (Cashikar *et al.*, 2014). Such flexibility is exceptionally highlighted by the unexpected discovery that the ESCRT-III’s IST1 and CHMP1B form a heteropolymer capable of scaffolding tubules with positive membrane curvature (McCullough *et al.*, 2015).

*In vitro* and *in vivo* analyses predominantly of the endocytic ESCRT arm have defined a step-wise activation of Snf7 by the sequential binding of the ESCRT-II, Vps25, to the ESCRT-III Vps20, which in turn releases the autoinhibition of Snf7 to induce filament formation (Teis *et al.*, 2008; Im *et al.*, 2009; Saksena *et al.*, 2009; Teis *et al.*, 2010; Henne *et al.*, 2012). Bro1-domain containing proteins like ALIX are also capable of activating Snf7 polymerization although through a distinct Snf7-binding interface (McCullough *et al.*, 2008). The ESCRT-III’s Vps2 and Vps24 are thought to alter the Snf7 filament helicity (Henne *et al.*, 2012) and recruit the AAA-ATPase Vps4, which stimulates ESCRT-recycling in a manner that might directly contribute to membrane scission (Teis *et al.*, 2008; Saksena *et al.*, 2009; Adell *et al.*, 2014).

Interestingly, Vps25 and Vps20 were absent from the genetic and functional analysis of ESCRT-III-mediated NPC assembly quality control, raising the possibility that Snf7 activation at the NE might require specific factors distinct from those at endosomes (Webster *et al.*, 2014). As the recruitment of ESCRT-III’s to different cellular compartments requires site-specific adaptor molecules (McCullough *et al.*, 2013; Hurley, 2015), it is possible that the adaptors themselves might contribute to the activation mechanism. This concept might be particularly relevant at the NE as the ‘orphan’ ESCRT, CHMP7 (Horii *et al.*, 2006), was recently shown to help recruit other ESCRT-III’s to seal NE holes at the end of mitosis (Vietri *et al.*, 2015). As CHMP7 in metazoans and in yeast (Chm7) might be a chimera of an ESCRT-II and ESCRT-III domain (Bauer *et al.*, 2015), a compelling hypothesis is that it might supplant the role of Vps25 and Vps20 at the NE.

The first identification of a role for ESCRTs at the NE was in budding yeast(Webster *et al.*, 2014)(aided by a genetic analysis in fission yeast (Frost *et al*., 2012)), which undergo a closed mitosis where the NE remains intact throughout the cell cycle. Snf7 and Vps4 were implicated in a surveillance pathway that ensures the assembly of functional NPCs, in part by preventing the formation of the “storage of improperly assembled nuclear pore complexes” (SINC) compartment, which is most prevalent in *vps4Δ* strains (Webster *et al.*, 2014). The SINC encompasses a NE subdomain where defective NPCs are aggregated during assembly and subsequently retained in mother cells to ensure daughter nuclear compartmentalization (Webster *et al.*, 2014). This study fits into a broader theme of emerging work supporting mechanisms that monitor NPC functionality and prevent the inheritance of defective NPCs (Colombi *et al.*, 2013; Makio *et al.*, 2013); the importance of understanding these mechanisms is reinforced by the observed loss of nups that occurs with age in rat brain neurons (D’Angelo *et al.*, 2009; Toyama *et al.*, 2013) and in budding yeast (Lord *et al.*, 2015).

Here, we further explore the mechanism of ESCRT-III function at the NE in budding yeast by focusing on the understudied ESCRT-II/III chimera, Chm7. Through a detailed analysis of the biochemical and genetic determinants of Chm7 localization at the NE, we uncover a previously undiscovered NE subdomain that requires Heh1 and expands upon NPC misassembly to form the SINC. Our data are consistent with the interpretation that the local regulation of Chm7 is required for NPC quality control.

## Results

### Heh2 directly binds to Chm7 and Snf7

We previously reported that Heh2 binds Snf7 through its N-terminal Lap2-emerin-MAN1 (LEM) containing domain, but it was not clear whether this interaction is direct (Webster *et al.*, 2014). To test this, we produced a recombinant heh2(1-308) that encompasses the entire extralumenal N-terminal domain of Heh2 (Figure 1A) and assessed binding to recombinant bead-bound GST-Snf7. We also tested binding to GST-snf7-N (Figure 1B), which encodes the ESCRT-III ‘core’ and lacks the acidic C-terminal auto-inhibitory helices thus mimicking its “open” active form. As shown in Figure 1C, we observed specific binding of heh2(1-308) and the GST-snf7-N, but not to full length Snf7. Despite the apparent specificity to the snf7-N construct over both GST and GST-Snf7, we were concerned that the sub-stoichiometric heh2(1-308) binding might reflect a weak interaction that could be dislodged by non-specific competition *in vivo*. However, we reproduced the direct (and specific) binding of heh2(1-308) to GST-snf7-N within an *in vitro* transcription translation mix where non-specific competitors are in abundance, suggesting this interaction is biologically relevant (Figure 1D). Further, by testing a series of truncations of the Heh2 N-terminal domain, we narrowed the binding interface between Heh2 and Snf7 to the N-terminal ~100 amino acids, which encompasses the LEM domain (Figure 1D); we were unable to test binding sufficiency with the LEM domain as we failed to produce a stable polypeptide. We nonetheless conclude that Heh2 directly binds to Snf7 thus providing a mechanistic basis for how this E-III is recruited to the NE. The mammalian orthologue of Chm7 helps seal NE holes at the end of an “open” mitosis (Vietri *et al.*, 2015) suggesting the existence of NE-specific adaptors that have not yet been defined capable of recruiting Chm7. We therefore tested whether Heh2 could directly interact with Chm7 as well. Interestingly, secondary structure and homology modeling support that Chm7 is a chimera of an ESCRT-II and ESCRT-III subunit, although the atomic structure remains to be solved (Figure 1‐figure supplement 1;(Bauer *et al.*, 2015)). Therefore, to test binding with heh2(1-308), we produced recombinant forms of Chm7 (Figure 1E) in addition to isolated ESCRT-II-(chm7-N) and ESCRT-III-like (chm7-C) domains. We also generated a C-terminal truncation that would model the “open” form of the potential ESCRT-III domain (chm7-CΔC). Remarkably, heh2(1-308) directly and specifically interacted with the Chm7 ESCRT-III domain in a manner analogous to its binding to Snf7, i.e. it bound to the ESCRT-III domain only in its putative “open” conformation (Figure 1F). These data reinforce the concept that the Chm7 C-terminal domain might have an ESCRT-III-like structure recognized by Heh2.

**Figure 1.**
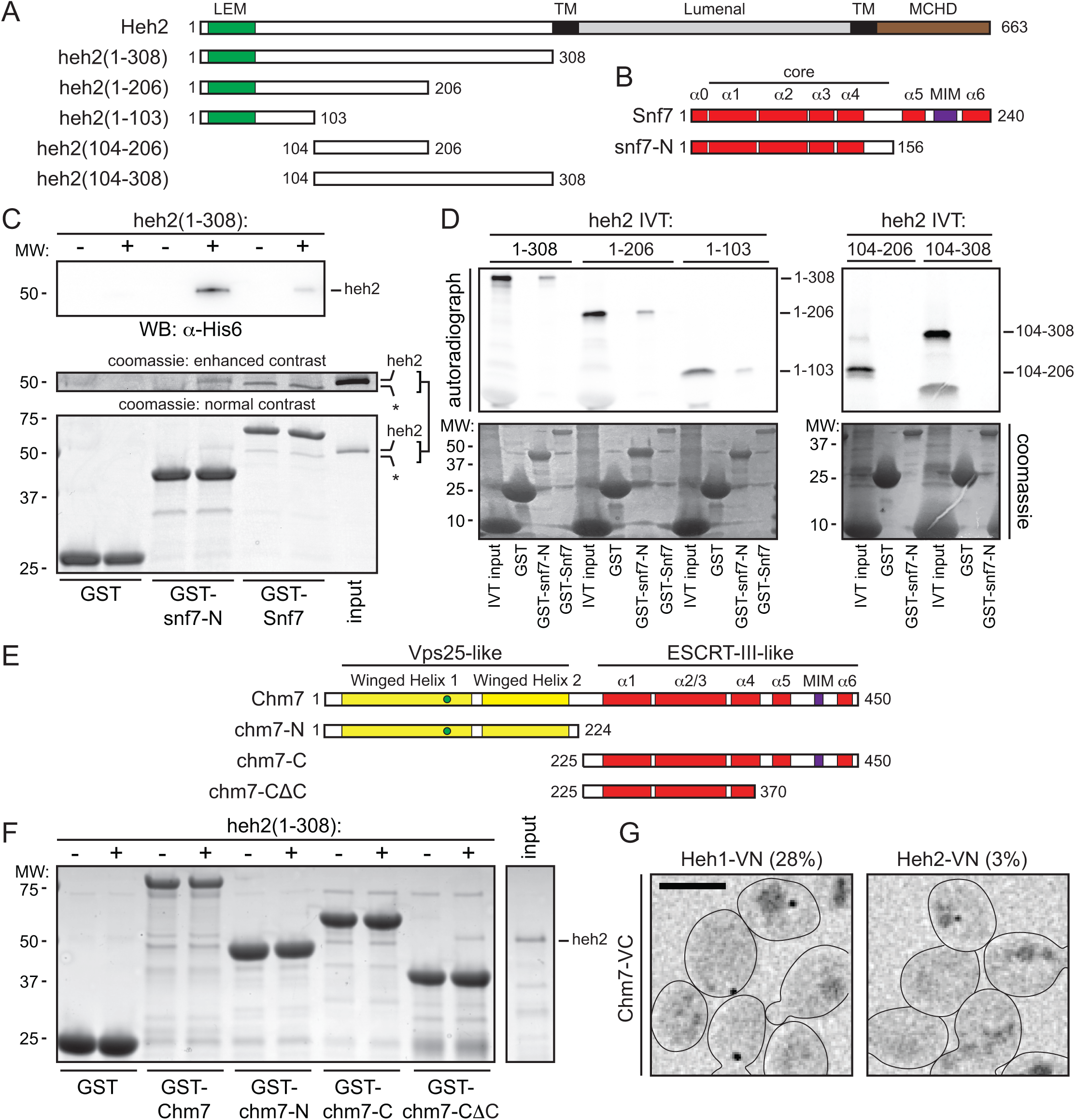
Heh2 directly binds Snf7 and Chm7. **(A, B)** Schematics of the domain organization and secondary structure of Heh2 and Snf7 truncations. LEM is Lap2-Emer-in-MAN1 domain (green), TM is transmembrane (black), MCHD is MAN1 C-terminal Homology Domain (brown), MIM is microtubule interacting motif (purple). Numbers are amino acid residues. **(C)** Recombinant purified GST, GST-Snf7 and GST-snf7-N were immobilized on GT-resin and incubated with buffer (-) or recombinant heh2(1-308)-His6. Bound proteins were eluted and separated by SDS-PAGE before detection by Western blot (top panel) or coomassie staining (bottom panel). Middle panel shows indicated cropped region of gel where contrast has been increased. Asterisk (*) marks GST-Snf7 degradation product. Numbers on side of gel show position of molecular weight (MW) markers. **(D)** In vitro transcription translation (IVT) reactions generating radiolabeled (S35) truncations of Heh2 (inputs). IVT reaction mixes were incubated with bead bound GST, GST-snf7-N or GST-Snf7 before washing, elution and detection of bound proteins by autoradiography (top panel) or coomassie staining (bottom). **(E)** Predicted secondary and tertiary structure of Chm7 (see also Figure 1 ‐ figure supplement 1) and generated truncations. **(F)** As in (C) except with GST-Chm7 constructs. SDS-PAGE gel is stained with coomassie. **(G)** BiFC experiments with yeast strains (BWCPL1816, 1817) expressing endogenously tagged Chm7-VC and either Heh1-VN or Heh2-VN. Scale bar is 5 μm. Deconvolved fluorescence micrographs shown are inverted. Percentages reflect the portion of population with detectable BiFC signal (n>100).

### Chm7 is required for Heh1/Heh2-Snf7 interactions *in vivo*

We next tested interactions between Chm7, other ESCRTs and Heh1/2 *in vivo* using Bifunctional Complementation (BiFC)(Figure 1G, Figure 1‐figure supplement 2) (Kerppola, 2008). Using this approach, bait and prey proteins are endogenously tagged with an N terminal (VN) and C-terminal (VC) domain of the fluorescent protein Venus, respectively. When bait and prey proteins associate, Venus folds and fluoresces. Consistent with the idea that Chm7 might bind directly to Snf7 (Bauer *et al.*, 2015), we observed specific BiFC between Snf7-VC and Chm7-VN within puncta throughout the cell but not between Bro1-VC or Vps20-VC and Chm7-VN (Figure 1‐figure supplement 2). We also observed low levels of BiFC between Chm7-VN and Vps4-VC likely mediated by the MIM domain in the Chm7 C-terminus ((Bauer *et al.*, 2015), Figure 1‐figure supplement 2). We were next able to recapitulate the direct binding interaction between Heh2 and Chm7 *in vivo* through the observation of BiFC signal between Heh2-VN and Chm7-VC as a solitary fluorescent focus (Figure 1G). Its low abundance (in only ~3% of cells) contrasted with cells expressing the Heh1-VN Chm7-VC combination where we observed a focus in ~28% of cells. This potential preference to Heh1 was not unanticipated, as Heh1 is a paralogue of Heh2 (King *et al.*, 2006) and it shares much of its primary and secondary structure including the N-terminal LEM domain; it is also established to have overlapping functional and physical interactions with Heh2 (Yewdell *et al.*, 2011; Webster *et al.*, 2014). Unfortunately, despite numerous attempts, we were unable to produce a stable recombinant form of the N-terminus of Heh1 to confirm its direct binding to Chm7. We nonetheless conclude that both Heh1 and Heh2 interact with Chm7.

Intriguingly, the Heh1/2-Chm7 BiFC fluorescence signal concentrated into one or two foci that were reminiscent of the NE-localized foci that we observed previously in BiFC experiments between Heh1/2 and Snf7 ((Webster *et al.*, 2014) and recapitulated in Figure 2B). We hypothesized that these BiFC foci might represent a functional organization of the ESCRT and Heh proteins at a discrete domain/region of the NE, although it is plausible that this pattern simply represents the product of stochastic molecular collisions stabilized by Venus folding. To differentiate between these possibilities, we first tested whether BiFC could readout dynamic functional ESCRT interactions at endosomes. For example, the functional interaction between Snf7 and Vps20 occurs after Vps25 action (Teis *et al.*, 2010). Consistent with these data, we failed to observe substantial BiFC signal between Vps20-VN and Snf7-VC in the absence of *VPS25* (Figure 2A,C), despite the fusion proteins being produced at levels comparable to wildtype cells (Figure 2‐figure supplement 1A). This effect was specific, as Snf7-VN Vps20-VC BiFC was not reduced in *vps24Δ, vps2Δ, bro1Δ, vps24Δ* and *chm7Δ* strains (Figure 2C). Thus, BiFC is capable of providing a faithful readout of functional spatiotemporal biochemical interactions of ESCRTs.

**Figure 2.**
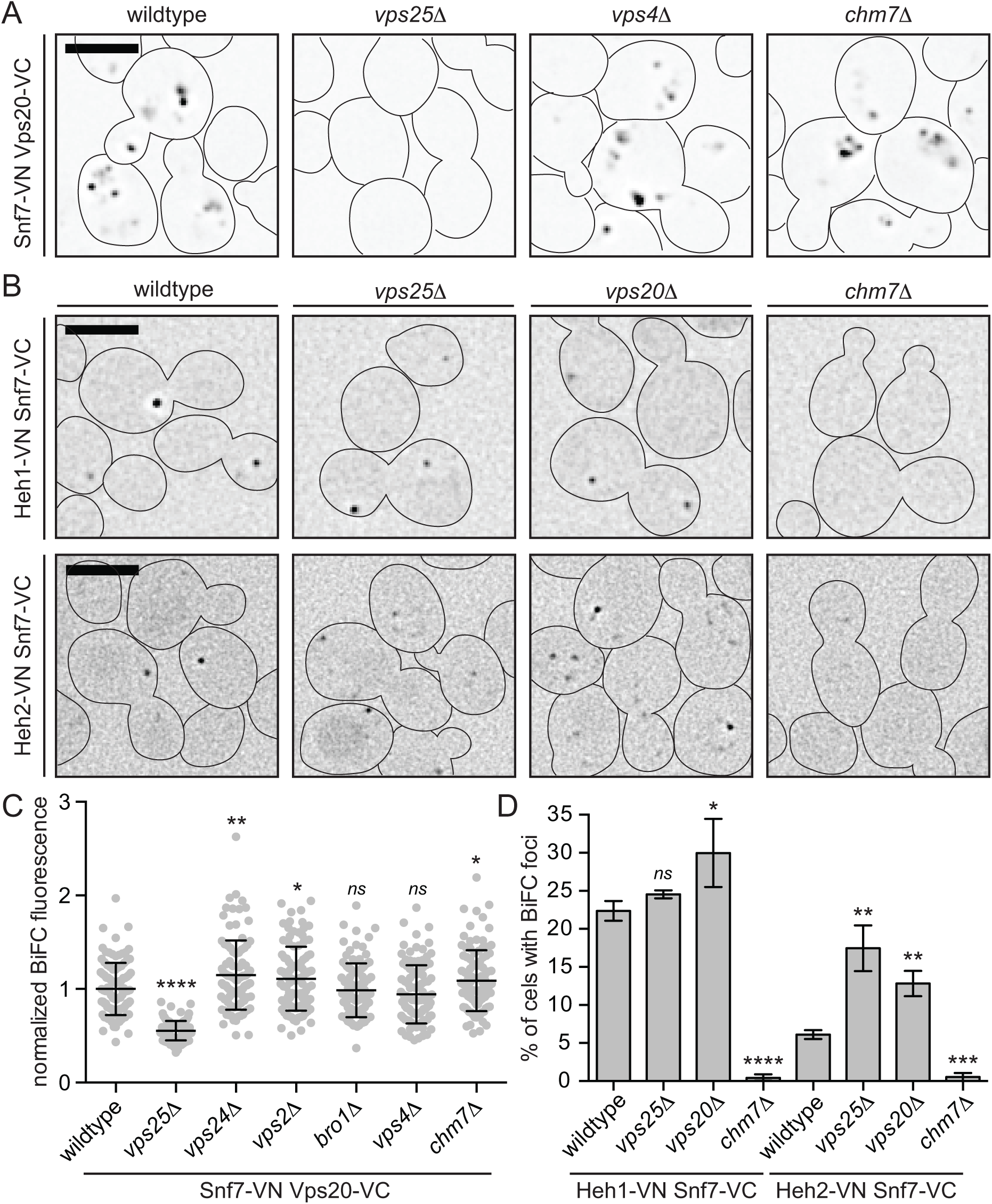
Chm7 is required for Heh1/2-Snf7 interactions *in vivo*. **(A, B)** BiFC of the indicated VN and VC fusions in the indicated strains (BWCPL1809-1815, SOCPL07, 13, 18-19, BWCPL1798-1800). All fluorescent images are deconvolved and inverted. Scale bar is 5 μm. See also Figure 2 ‐ figure supplement 1 for levels of VN and VC fusions within individual strains. **(C)** Plot of the total BiFC fluorescence between Snf7-VN and Vps20-VC within individual cells of the indicated strains. Data are from three independent replicates where 100 cells per strain were quantified. Error bars are standard deviation from the mean of each replicate. Statistical significance calculated using un-paired student’s T-test where ns represents p > 0.05; * is p ≤ 0.05; ** is p ≤ 0.01; **** is p ≤ 0.0001. **(D)** Plot of the percentage of cells with detectable BiFC of the Heh1-VN Snf7-VC and Heh2-VN Snf7-VC pairs. Data are from hree independent replicates where > 225 cells per strain per replicate were quantified. Error bars represent standard devia-ion from the mean of each replicate. Statistical significance calculated using un-paired student’s T-test where ns represents p > 0.05; * is p ≤ 0.05; ** is p ≤ 0.01; *** is p ≤ 0.001; **** is P ≤ 0.0001.

Knowing that BiFC could provide insight into the functional state of ESCRT-interactions, we tested whether deletion of different ESCRTs influenced the BiFC signal between Heh1/2 and Snf7. In contrast to BiFC at endosomes, and consistent with our prior genetic analysis (Webster *et al.*, 2014), the deletion of the canonical Snf7 activation components (Vps25 and Vps20) did not diminish, but in fact slightly increased BiFC of Heh1/2-VN Snf7-VC (Figure 2B,D). These data support the concept that there is a distinct biochemical arm of the ESCRT machinery at the NE; we therefore tested whether Chm7 might be a key component of this pathway by testing how it influenced Heh1/2-Snf7 BiFC. Strikingly, the deletion of *CHM7* virtually abolished BiFC between both Heh1/2-VN and Snf7-VC (Figure 2B,D) suggesting that Chm7 might help stimulate, stabilize and/or bridge the interaction between these proteins. In direct analogy to the Vps25-dependence of Vps20-Snf7 endosome interactions, these data also further reinforce the hypothesis that Chm7 (as a fusion of an ESCRT-II and ESCRT-III domain) might supplant the function of Vps25-20 at the NE. While the precise interplay between Heh1, Heh2, Chm7 and Snf7 remains to be fully understood within more extensive future biochemical and structural experiments, we think that it is likely that the localization of BiFC signal to a NE-focus represents a subdomain at the NE where these proteins interact to carry out specific NE-functions.

### Chm7 localizes to a focus on the NE

To begin to explore how Chm7 contributes to NE function, we examined its steady-state distribution by localizing Chm7-GFP expressed from the chromosomal *CHM7* locus (Figure 3A). Chm7-GFP showed a diffuse cytosolic fluorescence (consistent with recent work(Bauer *et al.*, 2015)) and, in ~26% of cells (Figure 4‐figure supplement 1C), a discrete focus (sometimes two foci; Figure 4B) that overlapped with a Nup170-mCherry NE marker (Figure 3A). As the Chm7-GFP NE focus resembled those observed within the BiFC experiments, we tested whether it enriched at a similar NE domain. As shown in Figure 3B, this is indeed the case as ~65% of the Heh1-VN Snf7-VC BiFC fluorescent foci co-localized with the Chm7-mCherry foci. As the appearance of the Chm7-GFP, and Heh1-Snf7 BiFC foci were highly reminiscent of SPBs, we tested whether they co-localized with the SPB protein, Mps3 (Figure 3B, Figure 3-figure supplement 1A,B). In both cases, we observed minimal co-localization, with only 14% of Chm7-GFP overlapping with Mps3-mCherry at steady state supporting the conclusion that the biochemical composition of this NE subdomain is distinct from SPBs.

**Figure 3.**
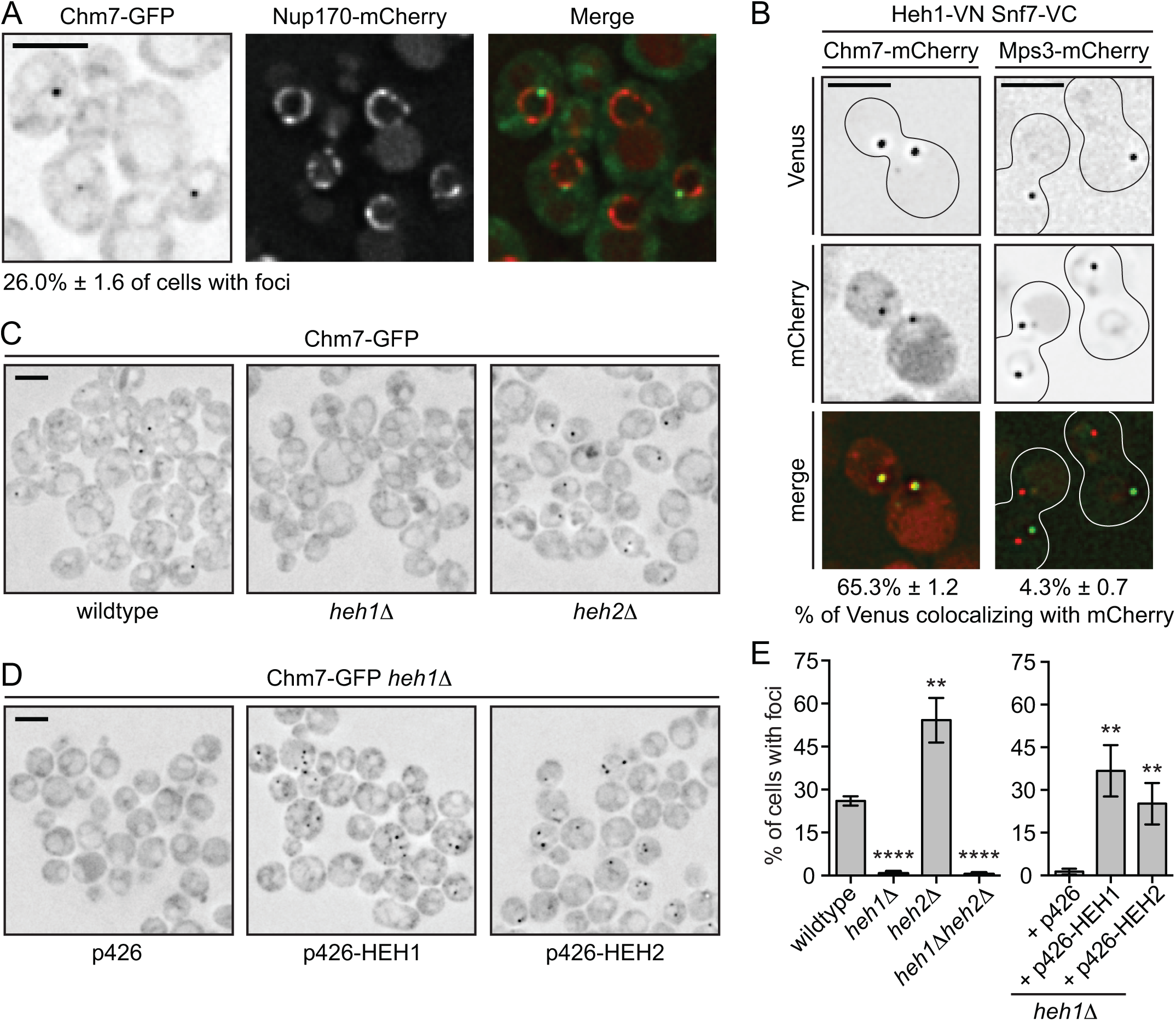
Chm7 localization to a NE subdomain requires Heh1. **(A)** Deconvolved fluorescent micrographs of BWCPL1840 expressing Chm7-GFP (left) and Nup170-mCherry (middle) with merged image (right). Only the green channel is inverted. Scale bar is 5 μm. Percentage of cells with Chm7-GFP NE-foci reflects the mean ± standard deviation derived from 3 independent replicates where > 200 cells were counted. **(B)** Deconvolved fluorescent micrographs of yeast strains expressing Heh1-VN and Snf7-VC with either Chm7-mCherry (BWCPL1842) or Mps3-mCherry (BWCPL1846). With the exception of the merge, all images are deconvolved inverted fluorescence micrographs. Scale bar is 5 μm. Percentages reflect mean ± standard deviation of the proportion of Chm7-or Mps3-mCherry foci that colocalize with Venus BiFC foci from 3 independent replicates where >40 Chm7-or >200 Mps3-mCherry foci were counted (See also Figure 3 – figure supplement 1B). **(C, D)** Deconvolved inverted fluorescence micrographs of the indicated strains (BWCPL1853-1855) expressing Chm7-GFP. In D, either an empty plasmid (p426) or those over-expressing *HEH1* or *HEH2* are introduced into the heh1Δ strain (BWCPL1853). **(E)** Plots of the percentage of cells from C and D with Chm7-GFP NE-foci. Data are from 3 independent replicates were >200 cells were counted for each strain. Error bars represent standard deviation from the mean. Statistical significance calculated using un-paired student’s T-test where ** represents a p value ≤ 0.01 and **** is p ≤ 0.0001.

### Heh1 contributes to the formation of the Chm7-subdomain

We next investigated how deletion of *HEH1* and *HEH2* influences the recruitment of Chm7-GFP to the NE. Strikingly, in *heh1Δ* cells, there was a complete loss of Chm7-GFP foci on the NE also seen in *heh1Δheh2Δ* cells (Figure 3C and E). In contrast, Chm7-GFP foci were more numerous and in a higher percentage of cells lacking *HEH2* (Figure 3C, E and Figure 4B); neither of these changes in distribution reflected alterations in Chm7 protein levels (Figure 4‐figure supplement 1A). However, consistent with the idea that both Heh1 and Heh2 can directly bind Chm7, the NE-recruitment of Chm7 could be restored in *heh1Δ* cells by either overexpressing *HEH1* or *HEH2* (Figure 3D,E). These data clearly suggest a requirement for Heh1 and Heh2 in the recruitment of Chm7 to the NE, but that other factors or physiological conditions influence the strength or stability of these interactions. To explore this relationship further, we used the distribution of Chm7-GFP at the NE as a facile assay to explore the molecular determinants of Chm7-GFP recruitment in more detail.

**Figure 4.**
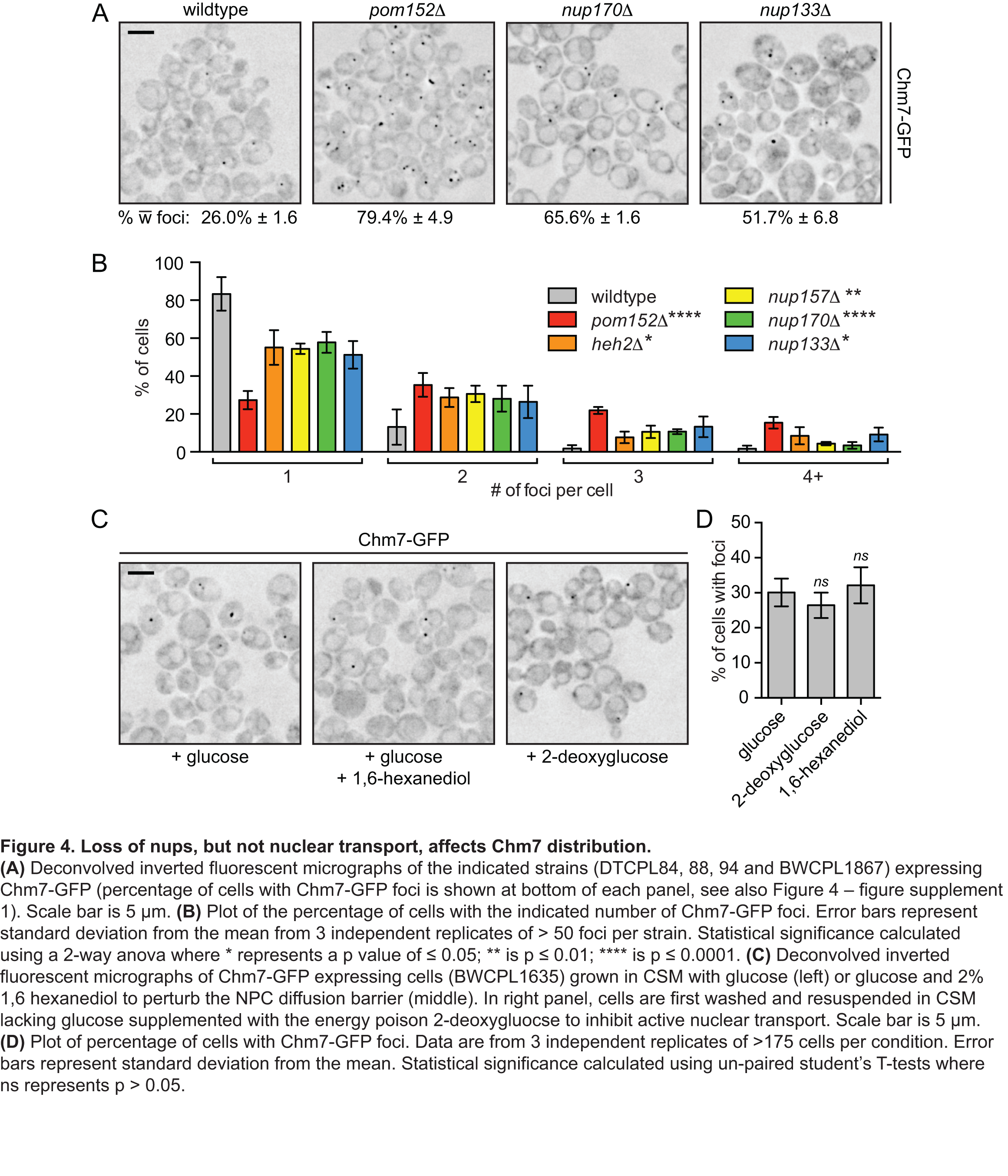
Loss of nups, but not nuclear transport, affects Chm7 distribution. **(A)** Deconvolved inverted fluorescent micrographs of the indicated strains (DTCPL84, 88, 94 and BWCPL1867) expressing Chm7-GFP (percentage of cells with Chm7-GFP foci is shown at bottom of each panel, see also Figure 4 – figure supplement 1). Scale bar is 5 μm. **(B)** Plot of the percentage of cells with the indicated number of Chm7-GFP foci. Error bars represent standard deviation from the mean from 3 independent replicates of > 50 foci per strain. Statistical significance calculated using a 2-way anova where * represents a p value of ≤ 0.05; ** is p ≤ 0.01; **** is p ≤ 0.0001. **(C)** Deconvolved inverted fluorescent micrographs of Chm7-GFP expressing cells (BWCPL1635) grown in CSM with glucose (left) or glucose and 2% 1,6 hexanediol to perturb the NPC diffusion barrier (middle). In right panel, cells are first washed and resuspended in CSM lacking glucose supplemented with the energy poison 2-deoxygluocse to inhibit active nuclear transport. Scale bar is 5 μm. **(D)** Plot of percentage of cells with Chm7-GFP foci. Data are from 3 independent replicates of >175 cells per condition. Error bars represent standard deviation from the mean. Statistical significance calculated using un-paired student’s T-tests where ns represents p > 0.05.

### Perturbation to NPC assembly, not nuclear transport, affects Chm7 distribution

As our prior work (Webster *et al.*, 2014) implicated the ESCRT machinery as contributing to normal NPC biogenesis, we next explored how Chm7-GFP localization was influenced by deletion of several non-essential nup genes, focusing on components of the membrane ring (Pom152), the inner ring (Nup170 and Nup157) and the outer ring (Nup133)(Figure 4-figure supplement 1B)(Alber *et al.*, 2007). Despite the fact that these nups are components of distinct NPC subcomplexes their perturbation results in a consistent increase in both the number of Chm7-GFP foci (Figure 4A, B) and the percentage of cells in which Chm7-GFP is localized to the NE at steady state (Figure 4‐figure supplement 1C), but not total Chm7-GFP levels (Figure 4‐figure supplement 1A). The most striking increase was observed in cells lacking *POM152*, which resulted in an increase from ~26% to ~80% of cells with Chm7-GFP NE foci, ~70% of which had more than one foci (Figure 4B). These data suggest that either the disruption of NPC function, or, perturbation in the assembly of NPCs lacking these components contribute to the recruitment of Chm7-GFP to the NE.

To differentiate between the possibilities that Chm7 recruitment to the NE responded to loss of NPC function or NPC assembly, we tested Chm7-GFP localization under conditions in which active nuclear transport is inhibited by incubating cells in the presence of the energy poison, 2-deoxyglucose (Shulga *et al.*, 1996; 2000; Timney *et al.*, 2006). We also treated cells with hexanediol, a common reagent used to perturb the diffusion barrier of the NPC (Ribbeck and Görlich, 2002; Shulga and Goldfarb, 2003). As shown in Figure 4C and D, neither of these treatments influenced the proportion of cells containing Chm7-GFP foci nor their relative numbers, suggesting that gross perturbation of NPC function does not influence Chm7 distribution.

We next examined Chm7-GFP localization upon blocking NPC assembly. For these experiments, we first assessed Chm7-GFP in *apq12Δ* strains, which are cold sensitive and show defects in NPC assembly and perturbations in NE morphology (Scarcelli *et al.*, 2007). Interestingly, even in wildtype cells the number of Chm7-foci was influenced by temperature changing from ~15% of the population at 23°C to ~50% at 37°C (Figure 4‐figure supplement 1C). These differences were exacerbated in *apq12Δ* cells, such that at lower (23°C) but most clearly at high (37°C) temperatures, there was a dramatic increase in the percentage of *apq12Δ* cells with foci (up to ~90%)(Figure 5A,B, Figure 5–figure supplement 1A). Perhaps most compellingly, ~37% of the *apq12Δ* cells had four or more foci at 37°C (Figure 5B). As *apq12Δ* cells also have aberrant NE morphology including NE herniations (Scarcelli *et al.*, 2007), we cannot be certain whether these alterations in Chm7 distribution reflect defects in NPC assembly or perturbations to membrane morphology (or both).

**Figure 5.**
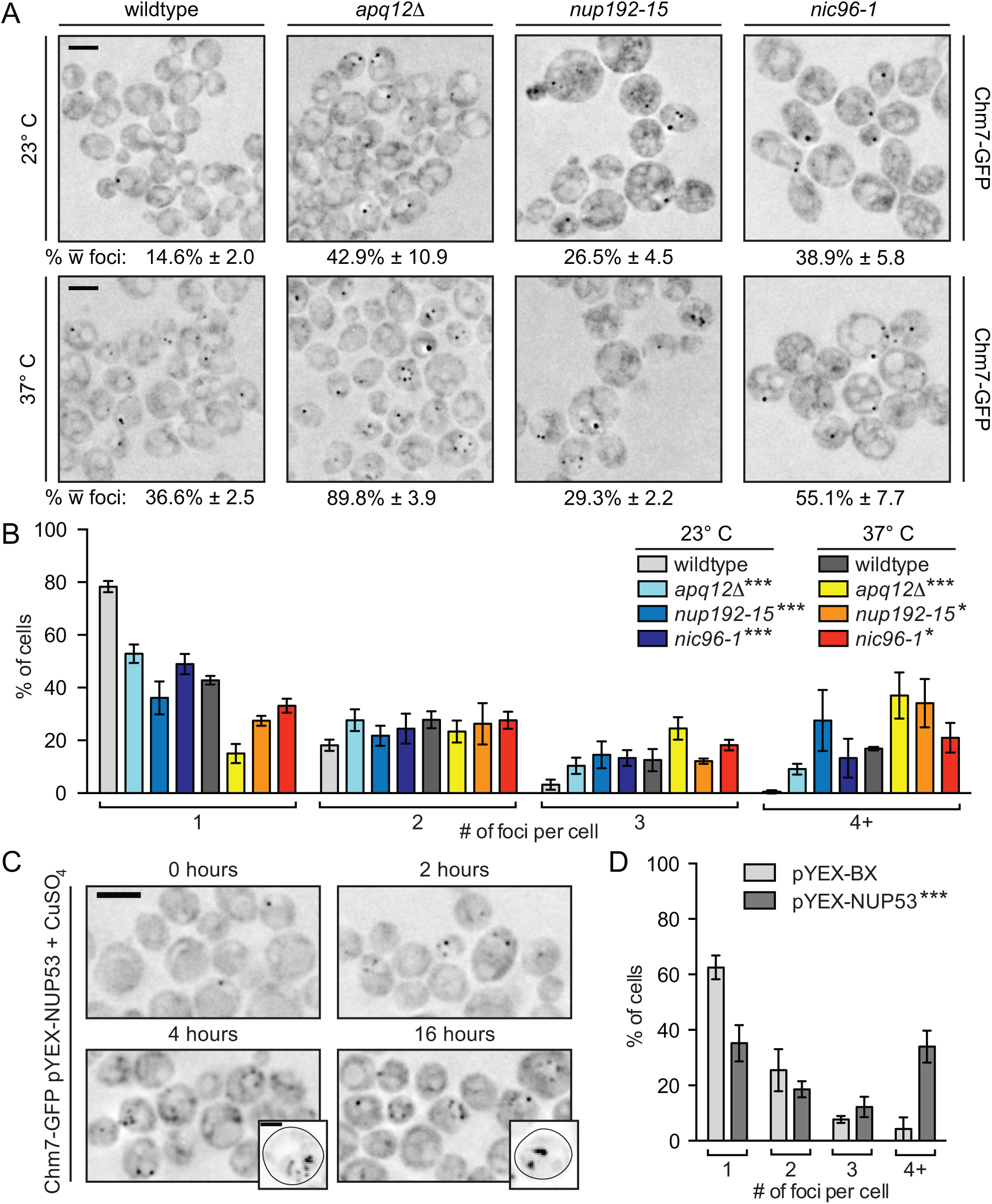
Chm7 is recruited to the NE when NPC assembly is blocked. **(A)** Deconvolved inverted fluorescence micrographs of strains expressing Chm7-GFP in the indicated temperature sensitive NPC assembly-deficient strains (BWCPL1840, DTCPL136, DTCPL228, DTCPL183). Scale bar is 5 μm. Top panels are at 23° C and bottom at 37° C. Percentages ± standard deviation of the proportion of cells where Chm7-GFP foci are visible are shown on bottom of each panel (see also Figure 5 – figure supplement 1). **(B)** Plot of percentage of cells from (A) with the indicated number of Chm7-GFP foci. Error bars represent standard deviation from the mean from 3 independent replicates of > 50 foci per strain. Statistical significance assessed with 2-way anova where * is p ≤ 0.05; ***, p ≤ 0.001. **(C)** Deconvolved inverted fluorescent images of Chm7-GFP expressing cells containing pYEX-NUP53, which expresses *NUP53* behind the copper-inducible *CUP1* promoter. Nup53 overproduction was induced by the addition of CuSO_4_ at time 0. Scale bar is 5 μm. Insets show maximum projection of cells where Chm7-GFP accumulates in patches reminiscent of intranuclear membranes induced by Nup53. Scale bar in inset is 2 μm. **(D)** Plot of the percentage of cells after 3 h of CuSO_4_ induction with the indicated number of Chm7-GFP foci. Error bars represent standard deviation from the mean from 3 independent replicates of > 50 foci per strain. Statistical significance assessed by 2-way anova where *** is p ≤ 0.001.

To begin to distinguish between whether the recruitment of Chm7 responded to changes in NE morphology and/or defects in NPC assembly, we examined Chm7 distribution within strains expressing the conditional nup alleles, *nic96-1* (Zabel *et al.*, 1996) and *nup192-15* (Kosova *et al.*, 1999). Both of these alleles encode components of the inner ring complex, which is essential for NPC assembly. Interestingly while at both the permissive (23oC) and non-permissive temperatures (37°C) there was a significant increase in the number of cells with Chm7-GFP (Figure 5‐figure supplement 1B) and a larger distribution in the number of foci per cell (Figure 5B), the most significant changes were observed at the lower temperature where these alleles are only partially compromised. Together, these data allow us to argue that it is likely that Chm7 is recruited to the NE when NPC assembly is perturbed, perhaps to defective NPCs or NPC assembly intermediates.

### Chm7 is recruited to lamellar membranes containing nuclear pores, and the SINC

To more directly assess whether Chm7 might interact with a site of NPC assembly (or misassembly), we took advantage of the observation that overexpression of Nup53 leads to formation of intranuclear lamellar membranes with pores devoid of NPCs but containing at least the transmembrane nups, Pom152 and Ndc1 (Marelli *etal*., 2001). Over 16 hours of Nup53 overproduction, we observed a consistent and incremental accumulation of Chm7-GFP onto these membranes (Figure 5C, D and Figure 5–figure supplement 1C, D). These data support that Chm7 is recruited to expanded regions of the NE known to have pore-like structures that lack fully formed NPCs. As these membranes were triggered by the extreme overproduction of Nup53, we also sought more physiological conditions to examine Chm7 distribution that would nonetheless mimic an environment rich in assembling, but malformed, NPCs.

Previously, we identified the SINC as a compartment on the NE containing newly synthesized nucleoporins, but not fully formed NPCs, which was most prevalent in *vps4Δ* and *vps4Δpom152Δ* cells (Webster *et al.*, 2014). We therefore asked whether Chm7 might enrich within the SINC. There was an obvious accumulation of multiple Chm7-GFP foci throughout both *vps4Δ* and, most substantially, *vps4Δpom152Δ* cells (Figure 6A). To visualize the SINC, we simultaneously evaluated the distribution of Nup170-mCherry (Webster *et al.*, 2014). As shown in Figure 6B, we observed a clear accumulation of Chm7-GFP in the SINC; indeed, virtually all SINCs contained Chm7 (Figure 6C). This contrasted with the lack of accumulation of Chm7-GFP within morphologically similar clustered NPCs observed in *nup133Δ* cells (Figure 6C, D). To quantify the level of Chm7 SINC accumulation, we measured the fluorescence intensity of Chm7-GFP in the SINC and compared these values to non-SINC Chm7-GFP foci that were similar to those seen in wildtype cells. Clearly, there was substantially more Chm7-GFP in SINCs reaching a 10-fold increase over non-SINC compartments (Figure 6E). Moreover, this SINC accumulation likely occurred over several generations as the levels of SINC-enriched Chm7-GFP correlated with the accumulation of Nup170- mCherry (Figure 6F). Consistent with this idea, SINC-associated Chm7-GFP is retained in mother cells during mitosis (Movie 1). Together, these data support that Chm7 can recognize a domain at the NE rich in assembling nups, but lacking fully formed NPCs. The enrichment of Chm7 in the SINC also suggests that the accumulation of Chm7 in the absence of Vps4 and Pom152 might contribute to the underlying molecular pathology that leads to the SINC’s expansion and resulting toxicity.

**Figure 6.**
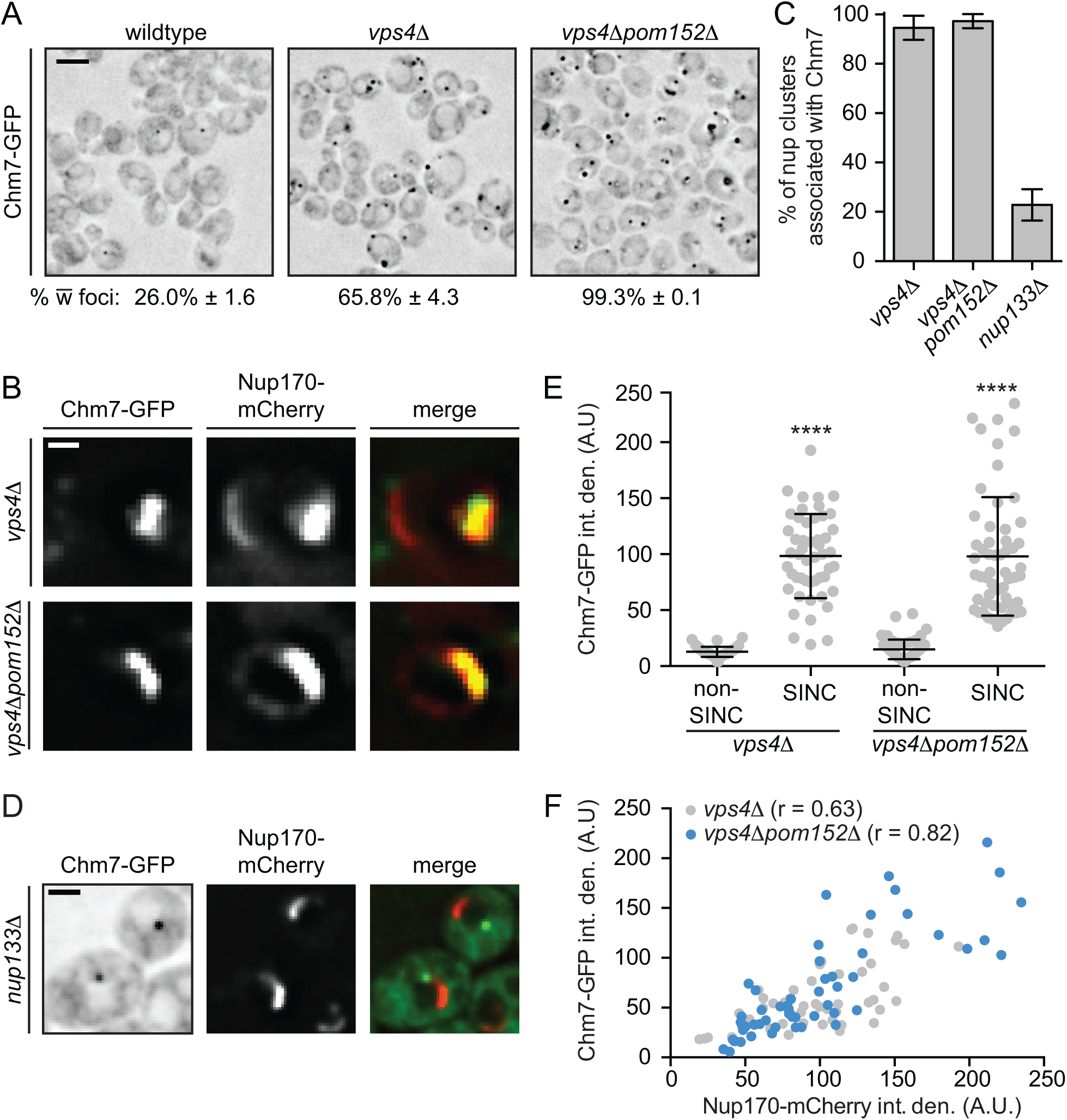
Chm7 accumulates in the SINC. **(A)** Deconvolved inverted fluorescence micrographs of Chm7-GFP in the indicated strains (DTCPL163, 131). Scale bar is 5 μm. The percentage of cells with Chm7-GFP foci is indicated below each panel (See also Figure 4 – figure supplement 1). **(B)** Representative deconvolved fluorescence images of SINC-containing nuclei in *vps4*Δ and *vps4*Δ*pom152*Δ cells expressing Chm7-GFP and Nup170-mCherry (green, red and merged images are shown). Scale bar is 1 μm. **(C)** Plot of the proportion of SINCs in *vps4*Δ and *vps4*Δ*pom152*Δ cells that colocalize with Chm7-GFP compared to its association with NPC clusters in *nup133*Δ cells (BWCPL1867). Error bars represent standard deviation of the mean where Chm7-GFP colocalization was assessed from > 50 NPC clusters in *nup133*Δ cells and 50 SINCs in *vps4*Δ and *vps4*Δ*pom152*Δ cells for 3 independent replicates. **(D)** Deconvolved fluorescent micrographs of BWCPL1867 expressing Chm7-GFP (left) and Nup170-mCherry (middle) with merged image (right). Only the green channel is inverted. Bar is 5 μm. **(E)** Plot of the total fluorescence (A.U.) of Chm7-GFP within the SINC compared to Chm7-GFP foci not associated with the SINC in *vps4*Δ and *vps4*Δ*pom152*Δ cells. Error bars represent standard deviation of the mean from > 50 SINC-associated Chm7-GFP or non-associated Chm7-GFP foci pooled from 3 independent replicates. Statistical significance calculated using un-paired student’s T-tests where **** represents p ≤ 0.0001. **(F)** Plot comparing the total fluorescence intensity of SINC enriched Chm7-GFP compared to Nup170-mCherry from individual SINCs in *vps4*Δ (DTCPL163) and *vps4*Δ*pom152*Δ (DTCPL131) cells. Linear regression calculated from 150 SINCs pooled from 3 independent replicates; r represents the linear correlation (Pearson’s) coefficient.

### Chm7 is required for SINC formation

To test the idea that Chm7 might be required to form the SINC in genetic backgrounds that promote its formation, we crossed *vps4Δpom152Δ* and *chm7Δ* cells and analyzed the growth of their progeny. Surprisingly, we observed a suppression of the impaired fitness of *vps4Δpom152Δ* strains with the *vps4Δpom152Δchm7Δ* strain showing growth indistinguishable from that of wildtype cells (Figure 7A). These data suggest that the growth retardation of *vps4Δpom152Δ* strains occurs due to a deleterious gain of function of Chm7. We considered two possible models to explain how deletion of Chm7 might rescue the *vps4Δpom152Δ* growth defect. In one, endocytic sorting defects in *vps4Δ* strains could be rescued by deletion of *CHM7*. As shown in Figure 7B and C, this did not seem to be the case, as the distribution of the model endocytic cargo Sna3-mCherry (Reggiori and Pelham, 2001; Russell *et al.*, 2012) remained in class E-compartments in both *vps4Δ* and *vps4Δchm7Δ* cells, and was not properly targeted to vacuoles as in WT and *chm7Δ* cells (Figure 7C). Alternatively, in the second model, Chm7 might directly contribute to SINC formation, which would then be the underlying cause of toxicity. Strikingly, and consistent with such a model, GFP-Nup49 no longer accumulated in the SINC in *vps4Δchm7Δ* or *vps4Δpom152Δchm7Δ* cells, despite its clear presence in *CHM7*-containing backgrounds (Figure 7D, Figure 7–figure supplement 1).

**Figure 7.**
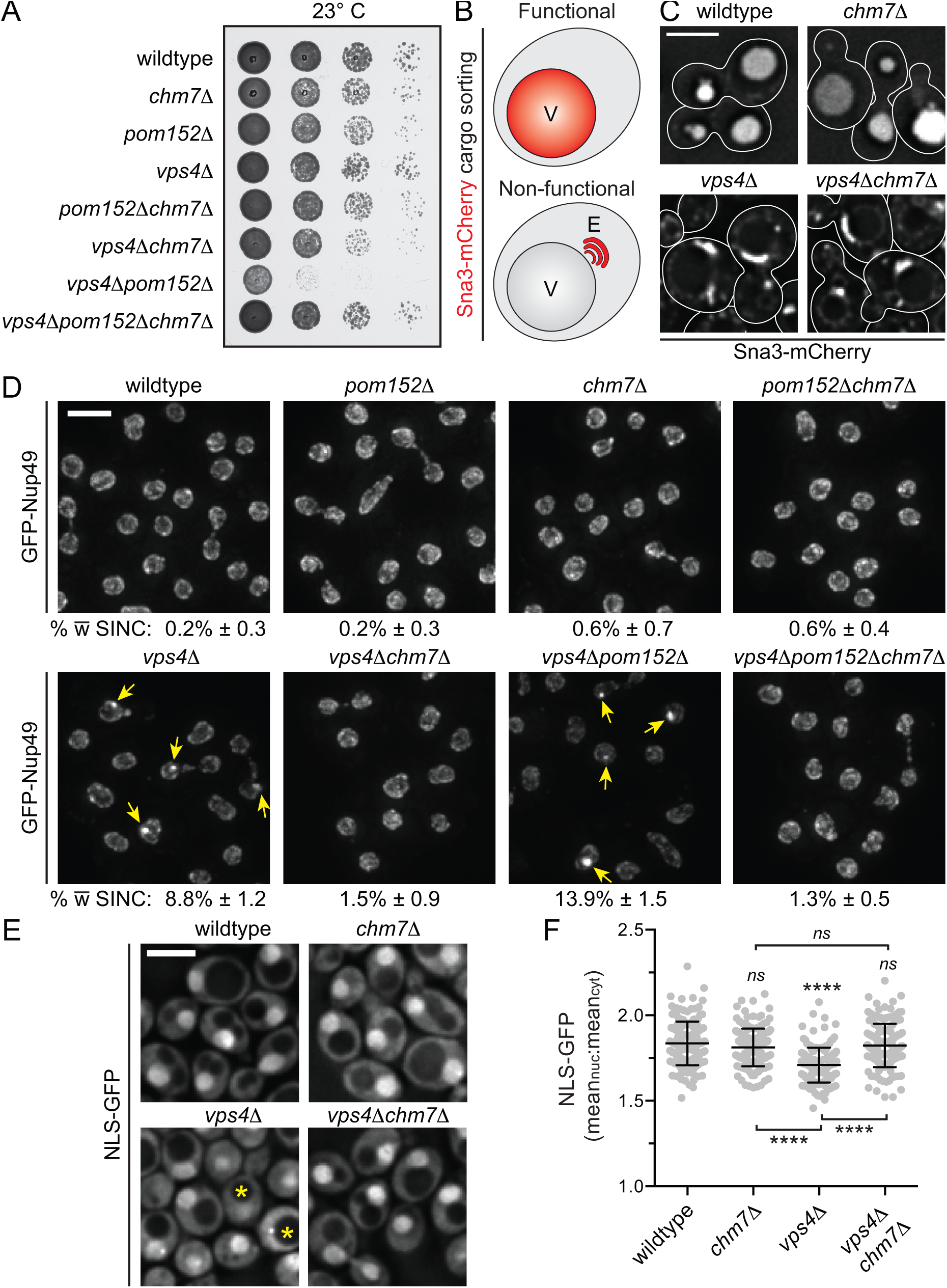
*CHM7* is required for SINC formation. **(A)** Deletion of *CHM7* rescues growth delays of *vps4*Δ*pom152*Δ cells. 10-fold serial dilutions of the indicated yeast strains (BWCPL1769-1775) grown on YPD at 23°C for 3 days. **(B)** Schematic of Sna3-mCherry localization. Sna3 accumulates in the vacuole (V) or class E compartment (E) in cells with, or lacking, ESCRT function, respectively. **(C)** Deconvolved fluorescence micrographs of Sna3-mCherry in the indicated genetic backgrounds (BWCPL1829-1832). Scale bar is 5 μm. **(D)** Deconvolved fluorescence micrographs of GFP-Nup49 in the indicated yeast strains. At bottom of each panel is the percentage of cells with SINCs (see also Figure 7 – figure supplement 1). Yellow arrows point out SINCs. **(E)** Deconvolved fluorescence micrographs of NLS-GFP in the indicated yeast strains (BWCPL1893-1896). The yellow asterisks point out cells with a loss of nuclear accumulation of the NLS-GFP reporter. Scale bar is 5 μm. **(F)** Plot of the mean nuclear to cytosolic ratio of NLS-GFP from (E). Error bars represent standard deviation from the mean from 3 independent replicates of 150 cells for each strain. Statistical significance calculated using un-paired student’s T-test where ns represents p > 0.05; ** is p ≤ 0.01; **** is p ≤ 0.0001.

The absence of the SINC in *vps4Δchm7Δ* cells also raised the possibility that the defects in nuclear compartmentalization observed in SINC-containing cells (Webster *et al.*, 2014) would be reversed in the absence of *CHM7*. To assess this possibility, we calculated the nuclear-cytosolic (N:C) ratio of a nuclear localization signal (NLS)-GFP reporter within individual cells, which had a mean value of 1.8 in wildtype and *chm7Δ* knockouts cell populations that was significantly reduced to 1.7 in *vps4Δ* strains (Figure 7E, F). Strikingly, N:C ratios in *vps4Δchm7Δ* were near-identical to wildtype cells (Figure 7E, F). Together, our data are consistent with a model in which the local regulation of Chm7 contributes to NPC quality control to promote proper nuclear compartmentalization.

## Discussion

NPC assembly occurs through the step-wise recruitment of nups and nup subcomplexes to an NPC assembly site concurrent with membrane remodeling events that lead to the fusion of the INM and ONM. The mechanisms coupling these two processes, nup recruitment and membrane remodeling, remain to be completely defined, but likely depend on the concerted action of many redundant factors. Indeed, in addition to specific nups and soluble nuclear transport receptors (Lusk *et al.*, 2002; Harel *et al.*, 2003; Ryan *et al.*, 2007; Lau *et al.*, 2009), there is a growing list of proteins that impact NPC assembly including established membrane-bending proteins with reticulon-domains (Dawson *et al.*, 2009; Chadrin *et al.*, 2010), factors that influence membrane fluidity like Apq12 (Scarcelli *et al.*, 2007) and Brr6 (Hodge *et al.*, 2010), integral INM proteins (Yewdell *et al.*, 2011; Talamas and Hetzer, 2011) and torsin (VanGompel *et al.*, 2015). Here, we introduce Chm7 as an additional component of the NPC assembly pathway; however, our data support that its role in nuclear pore biogenesis is distinct from these other established factors.

A critical clue to the function of Chm7 in the NPC assembly pathway is its specific recruitment to a solitary focus on the NE (Figure 3). With the exception of SPBs, which co-localize infrequently with Chm7, we are unaware of other proteins with this unique distribution, suggesting that Chm7 may delimit a previously undiscovered, biochemically distinct subdomain of the NE. While we do not yet have a complete accounting of the other factors that comprise this subdomain, the direct binding experiments (Figure 1), in concert with the genetic and BiFC analysis (Figure 2), support the conclusion that the integral INM proteins Heh1 and Heh2, the ESCRT-III subunit, Snf7, and Vps4 are key factors that contribute to its establishment and/or maintenance. Indeed, as the bulk of Heh1/2 are distributed throughout the NE (King *et al.*, 2006; Yewdell *et al.*, 2011), and Snf7 is found throughout the cytosol and at endosomes (Babst *et al.*, 1998), the localization of Chm7 provides a window into the function of this unique subset of NE-interactions; we propose that the Chm7 subdomain represents the major site of ESCRT-III function at the budding yeast NE.

Consistent with the concept that the Chm7 focus might demark the location of ESCRT-III function at the NE, the prevalence of the Chm7 foci dramatically increased upon inhibiting NPC assembly. Interestingly, however, the most abundant foci in *nic96-1* and *nup192-15* cells were present at the permissive (i.e. room temperature) instead of the non-permissive temperature (Figure 5). We interpret these data in a framework in which there are kinetic delays in assembly imposed by partially compromised alleles that is sensed and leads to Chm7 recruitment; these delays would be predicted to be absent at the non-permissive temperature due to a complete loss of function and assembly. This interpretation is consistent with the increase of foci in the non-essential nup knockouts strains as well (Figure 4), which allow for functional NPC formation but might nonetheless lead to a decrease in NPC assembly efficiency or quality.

Taken together, we suggest a model in which Chm7 is capable of recognizing a domain of the NE containing assembling (or misassembling) NPC(s). As very little is understood morphologically or biochemically about the steps in NPC assembly, it is difficult to assess at which “step” Chm7 might be recruited to a nascent NPC assembly site. An informative clue might be Chm7s enrichment on intranuclear lamellar membranes induced by the overexpression of Nup53 (Marelli *et al.*, 2001)(Figure 5). As these membranes are rich in NE pores, but not NPCs, a plausible hypothesis would be that Chm7 recognizes an NPC assembly intermediate post-membrane fusion. Such a model would also be compatible with the enrichment of Chm7 in the SINC, as the SINC contains FG-nups of the central transport channel (Webster *et al.*, 2014) that are likely assembled after fusion (Dawson *et al.*, 2009). Last, the concept that Chm7 might be recruited to a region of NE containing pores, but not NPCs, is consistent with work in mammalian cell lines where CHMP7 (and other ESCRT-III subunits) is recruited to seal NE holes at the end of mitosis by an annular fusion/membrane scission event (Vietri *et al.*, 2015; Olmos *et al.*, 2015), although how such structures are recognized remains unknown.

The scission of closely apposed membranes is the common thread that links ESCRT-III function in multiple cellular locales including the sealing of NE holes (Hurley, 2015). Such a mechanism, however, is not compatible with a role for ESCRTs in NPC assembly, as ESCRT-III would be predicted to seal the very pores necessary for their insertion. Thus, either Chm7 directly contributes to membrane remodeling to perhaps dilate or stabilize a nascent nuclear pore (Webster and Lusk, 2015), or, it is involved with a quality control pathway that prevents further aberrant assembly. One possibility would be the removal of defective NPCs or NPC assembly intermediates that would ultimately leave holes to be sealed by ESCRT-III. While NPCs are extremely stable in post-mitotic cells (D’Angelo *et al.*, 2009; Savas *et al.*, 2012; Toyama *et al.*, 2013), there is some evidence to support that NPCs might be removed from the NE in cell culture suggesting that this might be a plausible scenario (Dultz and Ellenberg, 2010). However, as *chm7Δ* cells do not have any obvious defects in nuclear compartmentalization (Figure 7) that would be predicted from the existence of empty NE holes, we favor an alternative quality control model where defective NPCs might be sealed off by expanded membrane.

The concept that defective NPCs might be sealed by exposed, closely apposed membranes can be found in decades-old electron micrographs of *nup116Δ* cells, in which NPCs are covered by a double membrane (Wente and Blobel,1993). Similar structures have been observed in *apq12Δ* cells (Scarcelli, *et al.*, 2007) and in those expressing *gle2* alleles (Murphy *et al.*, 1996) and could exist in the SINC as well (Webster *et al.*, 2014). As the seal is a double membrane, a hypothetical intermediate would be an expansion of an untethered nuclear pore membrane (Wente and Blobel, 1993); the sealing of such an expansion would be compatible with an ESCRT-driven scission step. This model makes the prediction that NPCs in *chm7Δnup116Δ* cells would fail to be sealed (an active topic of investigation in our lab). Moreover, this could be the mechanistic basis for the observed gain of function of Chm7 in *VPS4* and *POM152* null strains, which could lead to prematurely formed membrane seals over nascent NPCs thereby contributing to SINC formation. Thus, the spatial and/or temporal regulation of Chm7 by Vps4 and Pom152 at an NPC assembly site will be essential for ensuring it functions productively to maintain nuclear compartmentalization; our data support that Chm7 has the unique capability to function in a mechanism that differentiates functional and non-functional NPCs.

To ultimately elucidate the mechanism of Chm7 function in NPC quality control will require a better understanding of how it recognizes a nascent assembly site. In our prior study, we implicated Heh2 as a factor capable of differentiating between fully formed and nascent NPCs (Webster *et al.*, 2014). While Heh2 remains a key part of such a surveillance mechanism, the data presented here point to Heh1 as being the major factor recruiting Chm7 to the NE and thus acting before Heh2 (Figure 3). The fact that Chm7 is required for the Heh1/2-Snf7 *in vivo* interactions (Figure 2) is further suggestive of a stepwise mechanism where Chm7 might help bridge or stimulate interactions between Heh2 and Snf7, perhaps explaining their sub-stoichiometric binding *in vitro* (Figure 1). Lastly, a key part of the NPC surveillance mechanism will be in understanding the mechanism of Snf7 activation. Our data argue against the attractive model where Chm7 simply supplants Vps20-Vps25 function at the NE, as this model does not incorporate binding of Heh2 (and likely Heh1) to the ‘open’ forms of both Snf7 and Chm7 (Figure 1). Determining how the Heh proteins and Chm7 come together to activate a potentially unique Snf7 homo or heteropolymer will be an exciting feature of this emerging mechanism of NPC quality control that awaits future structural studies.

## Materials and Methods

### Yeast Strain Generation and Growth

All strains used in this study are listed in Supplementary file 1. Gene knockouts and fluorescent tagging of endogenous genes were performed by PCR-based integration using the pFA6a plasmid series (see Table 1: Plasmids, Supplementary file 2) as templates (Longtine *et al.*, 1998; Van Driessche *et al.*, 2005; Sung and Huh, 2007). Standard yeast protocols for transformation, mating, sporulation and tetrad dissection were performed as described in (Amberg, D. Burke and Strathern, 2005). Unless otherwise indicated, cells were grown to mid-log phase in YPAD (1% bacto yeast extract [BD] 2% bacto peptone [BD] and 2% D-glucose [Sigma] 0.025% adenine [Sigma]) or in complete synthetic medium (CSM) containing 2% D-glucose and lacking one or more amino acids at 30°C. For comparing relative growth rates of yeast strains, equivalent numbers of cells from overnight cultures were spotted in six, 10-fold serial dilutions onto YPD plates and imaged after 48 h at RT.

*APQ12* null cells (DTCPL136) were grown overnight at 30°C, divided and diluted into two cultures at an OD_600_ of 0.2. Individual cultures were then grown at 23°C or 37°C for 5 h before imaging. Similarly, strains expressing temperature sensitive NPC assembly alleles (*nic96-1* and *nup192-15*) were grown at the permissive temperature (23°C) and shifted to 37°C for 5 h before imaging.

To drive the overexpression of *NUP53* behind the copper inducible *CUP1* promoter, Chm7-GFP expressing cells were transformed with either pYEX-BX or pYEX-BX-NUP53. Resulting transformants were grown to mid-log phase in CSM-URA. CuSO_4_ was added to both cultures at a final concentration of 0.5 mM and imaged over 16 h (Marelli *et al.*, 2001).

### Plasmids

All plasmids are listed in Supplementary file 2. All plasmids generated in this study were verified by sequencing. To generate pRS426-HEH1 and pRS426-HEH2, the *HEH1* and *HEH2* genes were PCR amplified from isolated chromosomal DNA with 5’ and 3’ regions encompassing the native promoters and terminators. The PCR products were integrated into pRS426 (Christianson *et al.*, 1992) using homologous recombination in yeast. Briefly, pRS426 was linearized and co-transformed with the *HEH1* and *HEH2* PCR products into W303a. Transformants were selected on CSM-URA plates and grown overnight in CSM-URA before plasmid isolation using a modified protocol in (Amberg, D. Burke and Strathern, 2005) that incorporates a plasmid purification step on Qiagen miniprep columns (Qiagen).

All other plasmids were generated using the Gibson Assembly MasterMix (New England Biolabs) according to the manufacturer’s instructions. Briefly, coding regions of plasmid inserts were PCR amplified from W303 chromosomal DNA using Q5 DNA polymerase (New England Biolabs) and assembled into either pCS2 (Promega) linearized with BamHI/EcoRI (New England Biolabs), pGEX-6P1 (GE Life Sciences) linearized with BamHI/XhoI or pET28a linearized with XbaI/XhoI.

### Production and Affinity Purification of recombinant proteins

*E. coli* BL21 (DE3) cells expressing GST fusion proteins were grown overnight, diluted to an OD_600_ of 0.15 in 2xYT, and allowed to reach an OD_600_ of 0.5 before the addition of IPTG to a final concentration of 0.5 mM. Cultures were then shifted to 24°C for 6 h before collecting cells by centrifugation. Cell pellets derived from 50 mL of culture were flash frozen in liquid nitrogen and stored at −80°C before lysis and protein purification.

Frozen pellets were resuspended in 15 mL lysis buffer (50 mM Tris pH 7.4, 300 mM NaCl, 2 mM MgCl_2_, 2 mM CaCl_2_, 10 % glycerol, 0.5 % NP-40, 1 mM DTT, Roche complete protease inhibitors) and lysed by sonication (Branson Sonifier 450). Whole cel lysates were cleared by centrifugation for 15 minutes at 20000 × *g*. Supernatants were incubated in batch with 100 μL of glutathione sepharose (GT) beads 4B (GE Healthcare) for 1 h, collected by centrifugation, and washed three times with lysis buffer before being used as bait in binding experiments (below) or released from beads (and GST) through incubation with HRV 3C protease (Thermo Scientific) overnight at 4 °C.

### Recombinant protein binding experiments

GT-beads with bound GST, or GST-fusions of Snf7 or Chm7 were washed with a binding buffer of PBS containing 10 % glycerol, 0.5 % NP-40 and 0.5 mM DTT. Beads were then incubated with purified Heh2 generated from HRV 3C cleavage or *in vitro* transcription and translation (IVT; see below) for 1 h at 4 °C, collected by centrifugation and washed three times with binding buffer. Bound proteins were eluted with 30 μL of 2x SDS-PAGE sample buffer before separation by SDS-PAGE. Proteins were visualized by staining with SimplyBlue SafeStain (Invitrogen), Western blot or autoradiography.

### *In vitro* transcription and translation (IVT) and autoradiography

TNT Quick Coupled Transcription/Translation System (Promega) was used to generate radiolabeled fragments of Heh2. Specifically, individual 40 μL IVT reactions were performed for 90 minutes at 30°C and consisted of 600 ng of plasmid DNA encoding Heh2 fragments, 20 μCi [^35^S] Methionine and 10 uL of IVT mix. 10 μL of the completed IVT reaction was used within individual binding experiments. SDS-PAGE gels with radiolabeled proteins were dried at 80°C for 1 h using a gel dryer (EC355, E-C Apparatus Corporation) and exposed to autoradiography film (Bio-Rad Molecular Imager FX Imaging Screen) for three days. [^35^S] Methionine-labeled proteins were visualized using a Bio-Rad Personal Molecular Imager (Bio-Rad).

### Western blotting

Proteins separated by SDS-PAGE were transferred to nitrocellulose for Western blotting with the following primary antibodies: anti-GFP (gift of M. Rout), anti-Snf7 (gift of D. Katzmann). Primary antibodies were detected with anti-rabbit HRP conjugated secondary antibodies followed by ECL. ECL visualized using a VersaDoc Imaging System (Bio-Rad).

### Fluorescence microscopy

All fluorescence images were acquired on a DeltaVision widefield deconvolution microscope (Applied Precision/GE Healthcare) equipped with a 100x, 1.40 numerical aperture objective (Olympus), solid state illumination and an Evolve EMCCD camera (Photometrics). In all cases, Z-stacks of images (0.2 μm sections) were acquired. For timelapse experiments, cells were first immobilized on a 1.4% agarose pad containing CSM, 2% D-glucose, 0.025% adenine, and sealed with VALAP (1:1:1, vaseline:lanolin:paraffin) before imaging.

### Image processing and analysis

All fluorescent micrographs presented were deconvolved using the iterative algorithm in softWoRx (version 6.5.1; Applied Precision GE Healthcare) with subsequent processing and analyses performed in Fiji/ImageJ (Schindelin *et al.*, 2015). Importantly, quantification of fluorescence intensities was performed on unprocessed images after background subtraction.

To quantify the fluorescence intensity of Venus within BiFC experiments, the integrated density of Venus fluorescence within a central focal plane of an entire cell was measured. Similarly, the total fluorescence intensity of Chm7-GFP associated (or not) with SINCs was measured using Nup170-mCherry as a SINC landmark. To correlate the enrichment of Chm7-GFP and Nup170-mCherry in the SINC, the integrated density of Nup170-mCherry and Chm7-GFP fluorescence was measured and plotted on a correlation curve. The linear correlation coefficient (Pearson coefficient, r) was calculated using Prism (GraphPad).

To calculate the nuclear to cytosolic ratio of the NLS-GFP reporter, the integrated density of two identically sized regions of interest (one cytoplasmic, one nuclear) within a middle z-section of individual cells was measured and related.

### Deoxyglucose and hexanediol treatment

To disrupt the diffusion barrier of the NPC, cells were collected by centrifugation and resuspended in CSM containing 2% 1,6 hexanediol for 10 min. at RT before imaging (Shulga and Goldfarb, 2003). Similarly, to inhibit active nuclear transport, cells were collected by centrifugation and resuspended in CSM lacking glucose but supplemented with 10 mM 2-deoxy-D-glucose and incubated for 45 min. at 30°C before imaging(Shulga *et al.*, 1996).

### Protein alignment and structural modeling

The alignment of Chm7 and Vps25 in Figure 1 – figure supplement 1B was generated by Phyre2 (Kelley *et al.*, 2015). Phyre2 predicted and aligned the secondary (2°) structure of Chm7 to the 2° structure of Vps25 based on the crystal structure, PDB: 1xb4 (Wernimont and Weissenhorn, 2004) with a confidence of 96.6%. The tertiary (3°) structure of Vps25, which consists of two winged-helix domains (Teo *et al.*, 2004), were manually mapped onto the Vps25 sequence.

The alignment of Chm7, Snf7, and Vps20 was generated by T-Coffee Expresso alignment tools (Notredame *et al.*, 2000) in Figure 1 – figure supplement 1C. The illustration of the Chm7 2° structure is based on predictions generated by Phyre2 (Kelley *et al.*, 2015). Alpha helices 1-4 of the Snf7 2° structure correspond to the Snf7 “core” domain from the crystal structure, PDB: 5FD7a (Tang *et al.*, 2015). Phyre2 structural predictions of Chm7 showed high similarities between the Snf7 “core” domain and the corresponding Chm7 domains, which aligned with a confidence score of 97.9%. The remaining Snf7 2° structure/domain organization (alpha helix 0, 5-6, linker, and MIM domain) is based on predictions from (Tang *et al.*, 2015) and (Henne *et al.*, 2012). The outlined MIM domains of Chm7 and Vps20 are from (Bauer *et al.*, 2015) and (Shestakova *et al.*, 2010), respectively.

### Plots and statistical analysis

All graphs and statistical analyses were generated and performed using Prism (GraphPad 6). P-values for all graphs are represented as follows: ns, P > 0.05; * P ≤ 0.05; **, P ≤ 0.01; ***, P ≤ 0.001; ****, P ≤ 0.0001. All error bars represent standard deviation from the mean. For graphs in Figure 1 – supplement figure 2B, Figure 2C, Figure 6D, and Figure 7E, each circle represents one data point and means are marked by horizontal bars. Un-paired student’s T-tests were used to determine statistical significance for graphs in Figure 2C and D, Figure 3E, Figure 4D, Figure 4 – supplement figure 1C, Figure 5 – supplement figure 1A, B, and D, Figure 6D, Figure 7F, and Figure 7 – supplement figure 1. Two-way ANOVA tests using repeated measures by both factors were used to determine statistical significance for graphs in Figure 4B, Figure 5B and D.

## Acknowledgments

We are grateful for discussions of unpublished data with Adam Frost and Jeremy Carlton; we thank Megan King and members of the Lusk lab for critical reading of the manuscript. We appreciate the generosity of Topher Carroll, Mike Rout, Rick Wozniak and David Katzmann in sharing reagents and expertise. This work was supported by grants from the NIH GM105672 to C. Patrick Lusk. Brant Webster and David Thaller are also supported by NIH 5T32GM007223. Jens Jaeger is supported by the German Research Foundation (Deutsche Forschungsgemeinschaft, DFG).

---

**Figure 1 – figure supplement 1.**
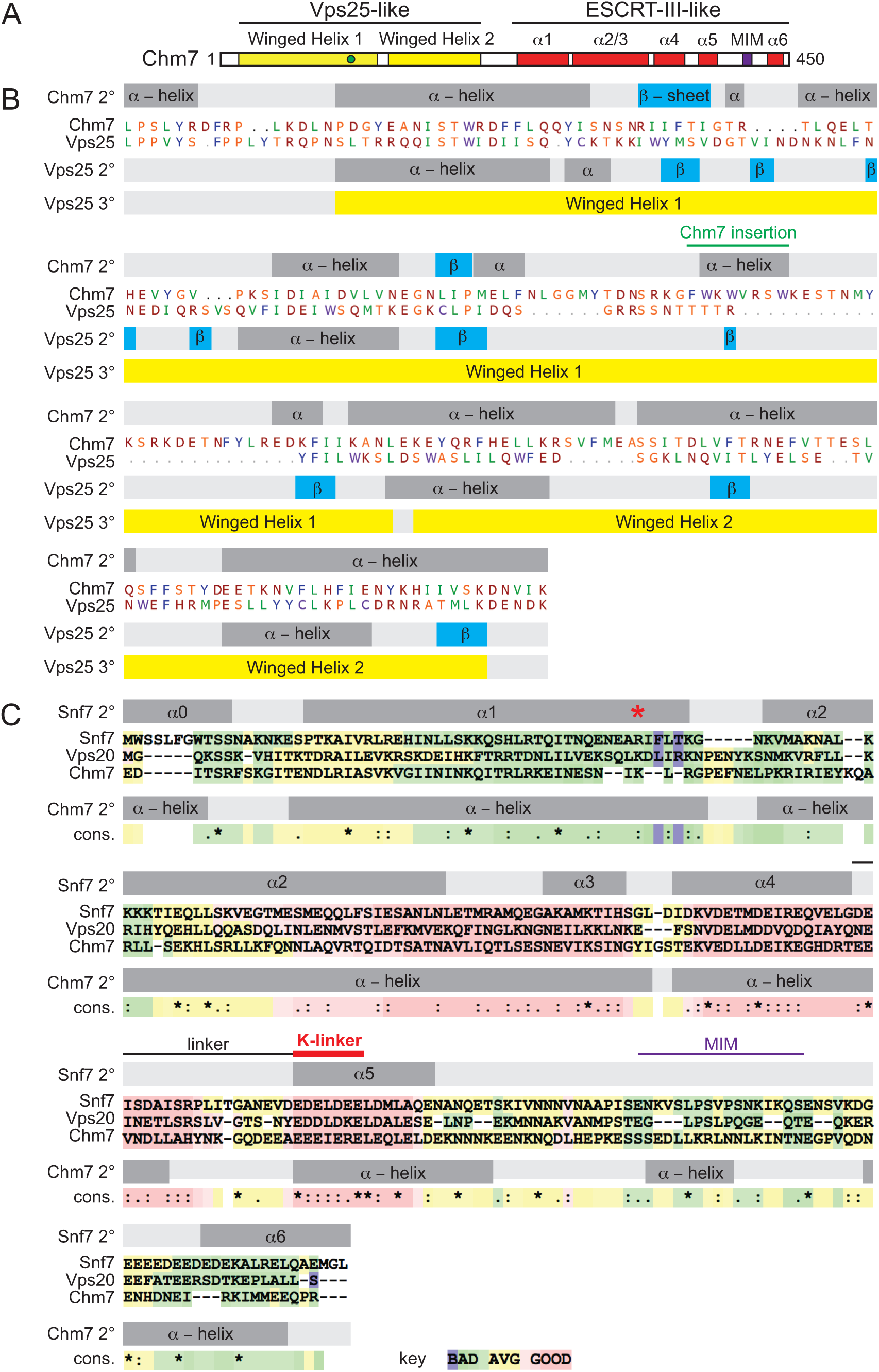
Chm7 is a chimera of ESCRT-II and ESCRT-III proteins. **(A)** Schematic of the predicted domain architecture and structure of Chm7. Green circle represents a Chm7-specific insertion (relative to Vps25) predicted by Phyre2. **(B)** Phyre2 generated alignment of the N-terminal domain of Chm7 (amino acids 10-214) and Vps25 (amino acids 4-195) with predicted secondary (2°) and tertiary (3°) structure. Alpha helices are gray; β-sheets are blue, winged helix domains are yellow. **(C)** T-Coffee Expresso generated alignment of the C-terminal domain of Chm7 (aa221-450) with Snf7 and Vps20. Coloration reflects the conservation score depicted in key at bottom. The Chm7 2° structural prediction was generated using Phyre2. Alpha helices are gray, red asterisk denotes a key residue for the intramolecular contact with alpha helix 5 and K-linker, K-linker is marked by red line, MIM domains marked by purple line.

**Figure 1 – figure supplement 2.**
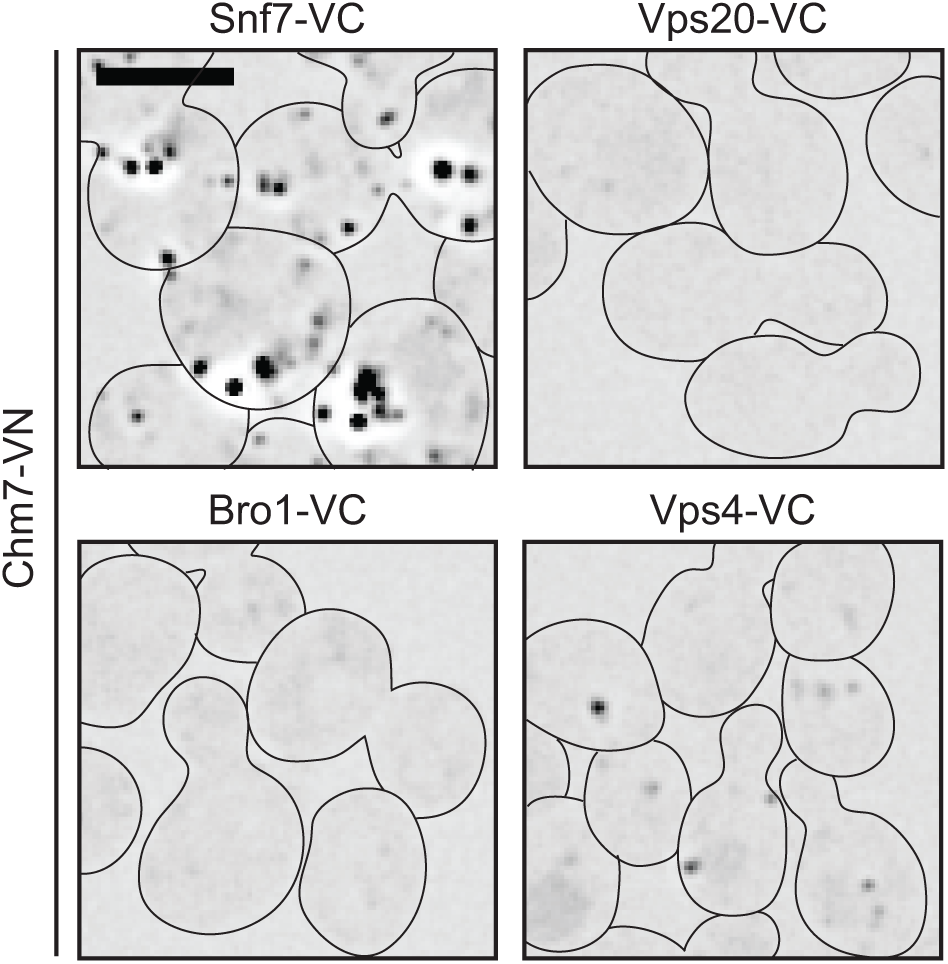
Chm7 interacts with Snf7 and Vps4 *in vivo*. BiFC experiments with strains (BWCPL1820-1823) expressing endogenously tagged Chm7-VN and either Snf7-VC, Vps20-VC, or Vps4-VC. Deconvolved fluorescence micrographs shown are inverted. Scale bar is 5 μm.

**Figure 2 – figure supplement 1.**
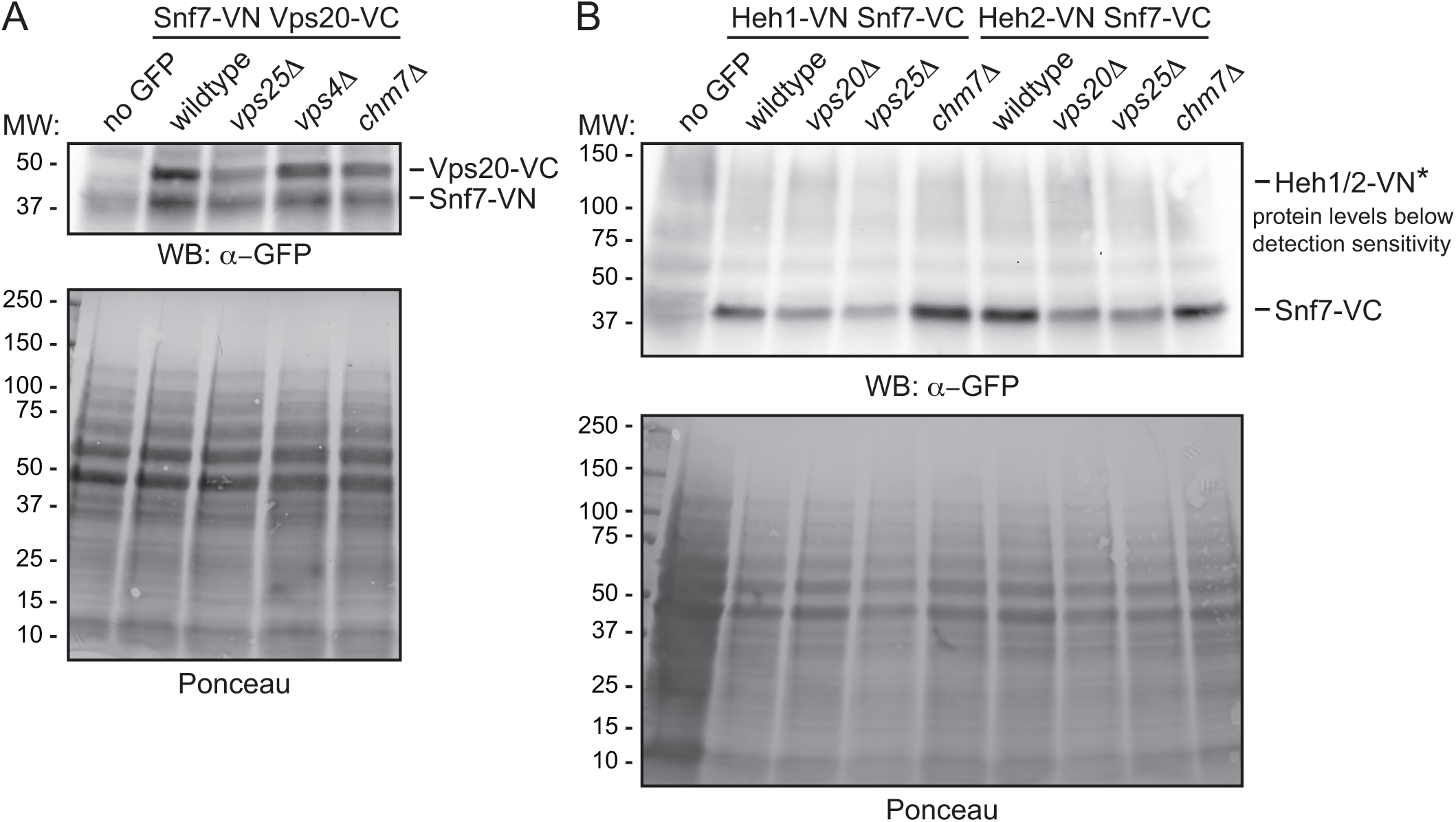
Split-venus fusion protein levels are unchanged in ESCRT null strains. **(A, B)** Western blots (WB) with polyclonal anti-GFP antibodies that can detect both VN and VC fusions in the indicated strains (A: BWCPL1809, 1810, 1814, and 1815; B: SOCPL7, 13, 18-19). To assess relative protein loads, bottom panel shows corresponding ponceau-stained nitrocellulose membranes.

**Figure 3 – figure supplement 1.**
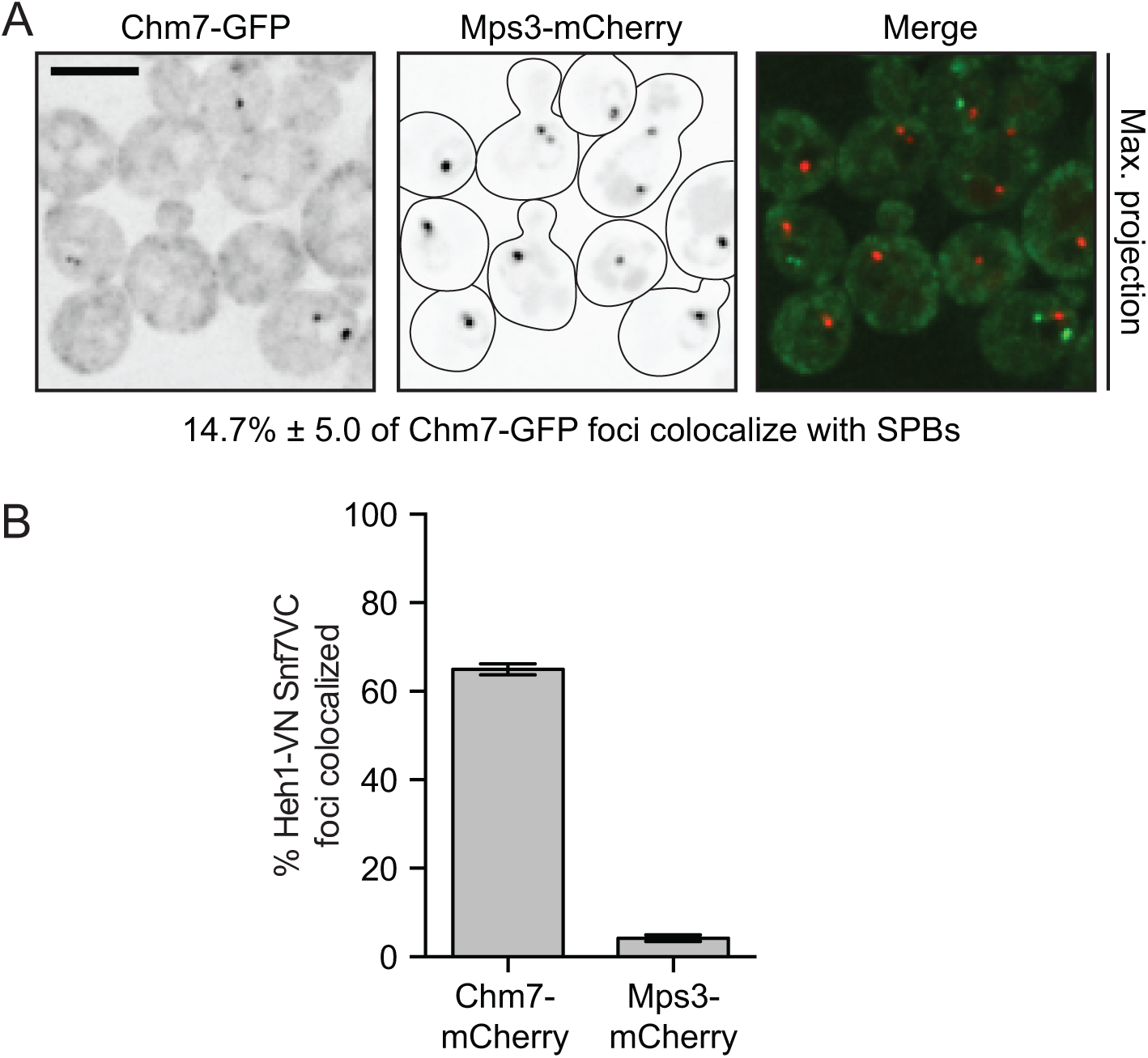
Chm7 does not associate with SPBs in the majority of cells at steady state. **(A)** A maximum intensity projection of deconvolved fluorescent micrographs of BWCPL1838 expressing Chm7-GFP (inverted, left) and Nup170-mCherry (inverted, middle) with merged image (right). Maximum projections shown to facilitate visualization of GFP and mCherry foci; colocalization was assessed from single z-planes. Scale bar is 5 μm. Percentage represents the mean ± standard deviation of the proportion of Chm7-GFP NE foci that colocalize with Mps3-labeled SPBs from 3 independent replicates where > 50 Chm7-GFP foci were counted. **(B)** Plot of the proportion of Heh1-VN Snf7-VC foci that colocalize with either Chm7-mCherry (BWCPL1842) or Mps3-mCherry (BWCPL1846). Data are from 3 independent replicates of > 50 Chm7-GFP foci (~150 cells). Bars represent standard deviation from the mean.

**Figure 4 – figure supplement 1.**
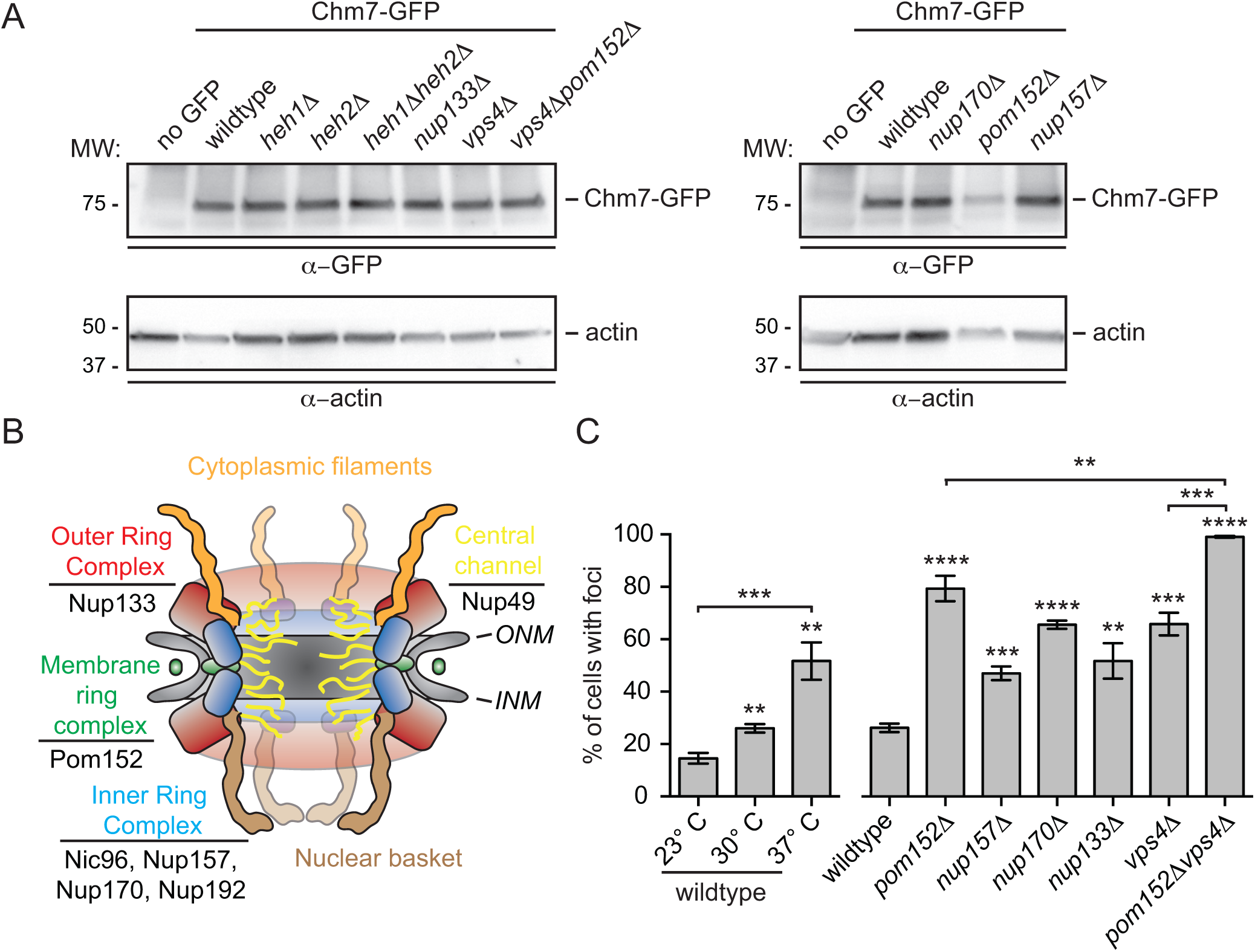
Chm7-GFP foci accumulate in nup knockouts. **(A)** Western blot of Chm7-GFP levels in the indicated strains (BWCPL1635, 1853-1855, 1867, DTCPL125, DTCPL131, DTCPL84, DTCPL88, DTCPL94) with actin loading control. **(B)** Schematic of the yeast NPC depicting nup subcomplexes (colors) and relevant subunits. Listed nups are referred to in text. **(C)** Plot of the percentage of cells with Chm7-GFP foci in the indicated strains (BWCPL1867, DTCPL125, DTCPL131, DTCPL84, DTCPL88, DTCPL94, DTCPL136). Error bars represent standard deviation from the mean from 3 independent experiments where > 175 cells per strain or condition were assessed. Statistical significance calculated using un-paired student’s T-test where ** represents p ≤ 0.01; *** is p ≤ 0.001; **** is p ≤ 0.0001.

**Figure 5 – figure supplement 1.**
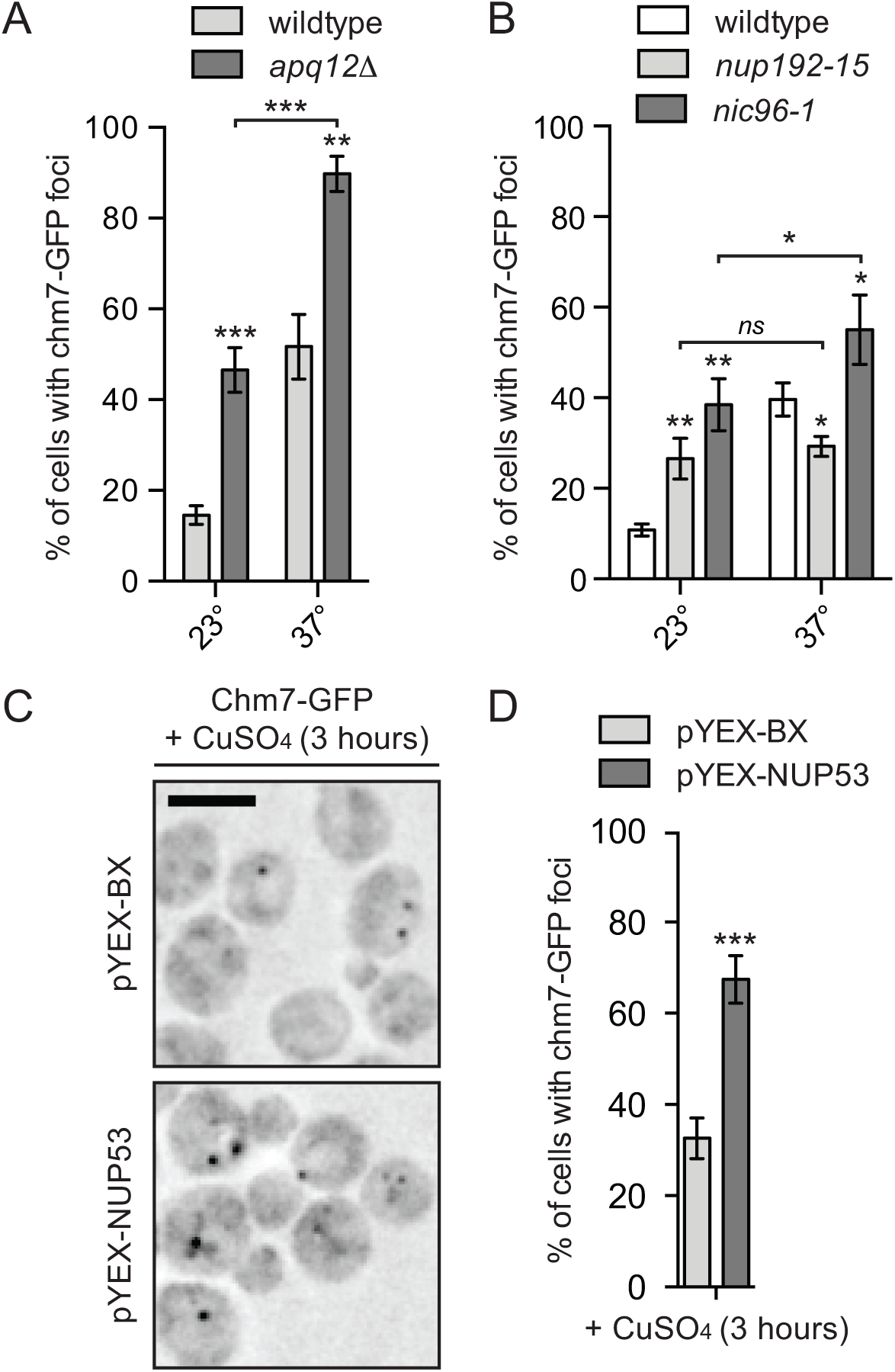
Chm7-GFP foci analysis in NPC assembly mutants. **(A,B, D)** Plot of the percentage of cells with Chm7-GFP foci in the indicated proportion of cells (DTCPL136, 228, 183, and BWCPL1635). Error bars represent standard deviation from the mean from 3 experiments of > 175 cells per strain or condition. Statistical significance calculated using un-paired student’s T-tests where ns represents p > 0.05; * is p ≤ 0.05; ** is p ≤ 0.01; *** is p ≤ 0.001. **(C)** Deconvolved inverted fluorescence micrographs of Chm7-GFP transformed with pYEX-BX or pYEX-BX-NUP53 in the presence of copper for 3 h. Scale bar is 5 μm.

**Figure 7 – figure supplement 1.**
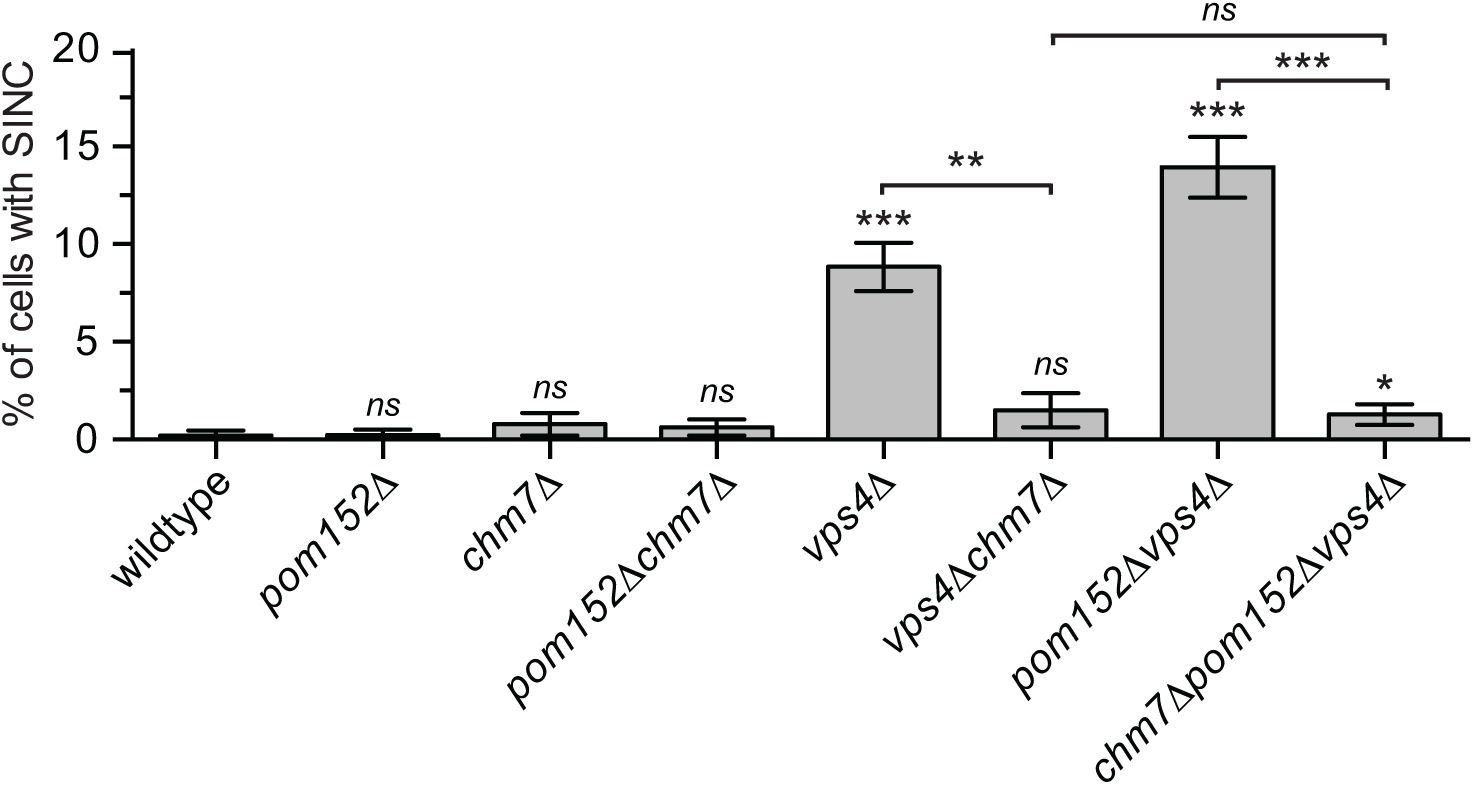
Deletion of *CHM7* suppresses SINC formation. Plot of proportion of cells in the indicated strains with SINCs. Error bars represent the standard deviation from the mean from 3 independent replicates quantifying >300 cells for each strain. Statistical significance calculated using un-paired student’s T-tests where ns is p > 0.05; * is p ≤ 0.05; ** is p ≤ 0.01; *** is p ≤ 0.001.

**Supplementary file 1.**
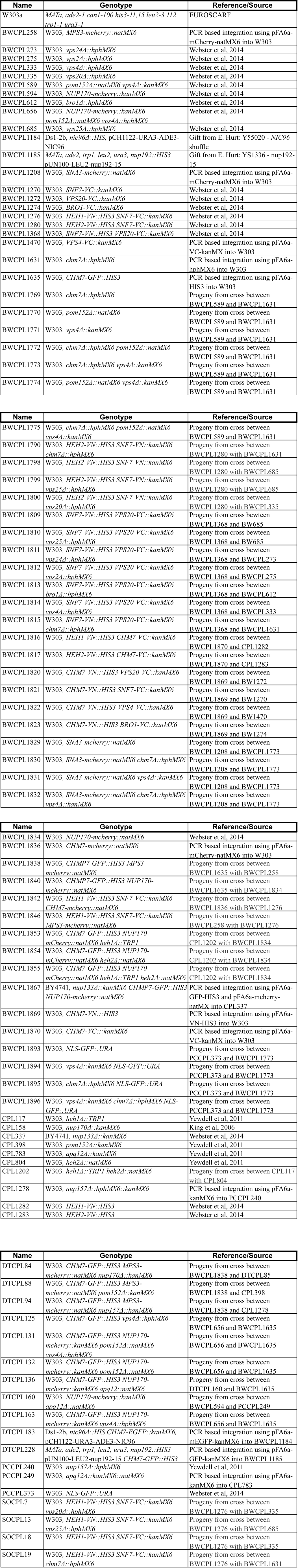
Yeast strains table

**Supplementary file 2.**
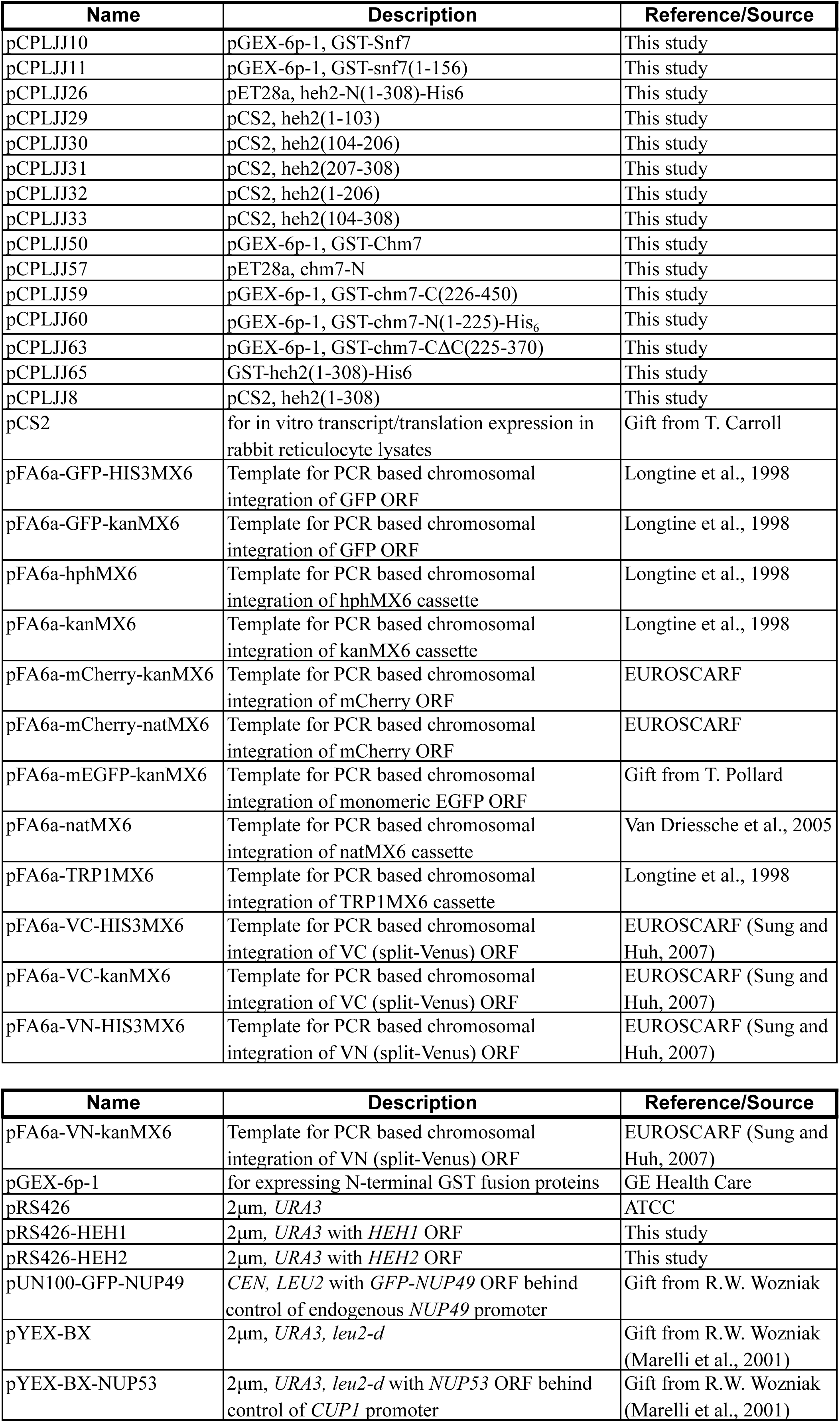
Plasmids Table

